# A luminal proteome of the endoplasmic reticulum and Golgi apparatus reveals a novel modulator of ER stress tolerance in African trypanosomes

**DOI:** 10.64898/2026.04.03.716285

**Authors:** Siqi Shen, Farnaz Zahedifard, Emmanuel Ayodeji Agbebi, Anna Zavrelova, Johanna Krenzer, Carla Gilabert Carbajo, Susanne Kramer, Calvin Tiengwe, Martin Zoltner

## Abstract

African trypanosomes employ specialised mechanisms of membrane trafficking as a key strategy to persist in both the mammalian host and insect vector. Their survival and pathogenicity rely on the continuous synthesis and surface delivery of extremely abundant surface coat proteins, imposing an extraordinary biosynthetic burden on the secretory pathway. Despite this, the luminal proteome of the *T. brucei* endoplasmic reticulum (ER) and Golgi apparatus remains incompletely characterised. Here, we exploit TurboID proximity biotinylation, using the abundant ER chaperone BiP (Binding-immunoglobulin protein) as luminal bait to map the ER proteome in bloodstream and procyclic lifecycle stages of *Trypanosoma brucei*. Comparison with BiPN, a truncated secretory form of BiP that transits the Golgi, provides differential compartmental labelling, together identifying 366 (BiP) and 428 (BiPN) proximity partners respectively and encompassing established ER quality control machinery, secretory cargo, and Golgi proteins. Quantitative ranking of BiP labelling intensity identifies a cohort of candidate BiP interactors: the most strongly enriched is Tb927.5.1160, a protein sharing structural homology with the mammalian BiP nucleotide exchange inhibitor MANF (mesencephalic astrocyte-derived neurotrophic factor). Endogenous mNeonGreen tagging confirms ER localisation of TbMANF in both life cycle stages, and reciprocal manipulation of its abundance by RNAi and inducible expression produces opposing shifts in cellular sensitivity to ER stress. These data are consistent with a role in regulating BiP ATPase cycling in an organism that, unlike yeast and mammals, lacks a canonical unfolded protein response, making TbMANF the first candidate regulator of BiP activity identified in kinetoplastids. Finally, TurboID proximity labelling anchored at the inner face of the nuclear pore via NUP65 extends our endomembrane map to the inner nuclear membrane, identifying candidate proteins of this specialised ER-continuous domain.

**AUTHOR SUMMARY:** African sleeping sickness is caused by *Trypanosoma brucei*, a parasite that survives in the mammalian bloodstream by constantly renewing its protective protein coat. To synthesise and export this surface coat, the parasite relies on two intracellular compartments, the endoplasmic reticulum (ER) and the Golgi apparatus, which function as a quality control and sorting factory for proteins entering the secretory pathway. However, the identity of the proteins that populate these compartments in blood-stage parasites, and that maintain functioning under stress conditions, has remained poorly mapped. Here, we used an enzyme-based proximity labelling strategy that identifies neighbouring proteins in live cells without disturbing their targeting signals, generating a comprehensive protein inventory of both compartments across the two main *T. brucei* lifecycle stages. Among the most strongly labelled proteins was Tb927.5.1160, a protein structurally related to a mammalian regulator of the master ER chaperone BiP. Reducing or increasing the abundance of Tb927.5.1160 in parasites produced opposite changes in ER stress tolerance, identifying it as a candidate modulator of ER homeostasis in a lineage that regulates protein quality control through mechanisms distinct from those operating in yeast or human cells. Together, our findings provide a new molecular resource for understanding how *T. brucei* sustains secretory pathway function under the biosynthetic demands of mammalian and insect host infection.

## INTRODUCTION

The classical secretory pathway is initiated in the ER, where nascent proteins destined for secretion encounter specialised conditions tailored for their folding, maturation and posttranslational modification. Secretory cargo undergoes various quality control steps and is finally passed to the *cis*-Golgi, from where it is trafficked across the stacks of Golgi cisternae to the *trans*-Golgi network (TGN). Protein glycosylation, initiated in the ER, is continued and elaborated in the Golgi apparatus where glycosyltransferases sequentially elongate the carbohydrate chains [1]. The TGN functions as a sorting hub for protein cargo routed to the plasma membrane, the endosomal compartments and other organelles. Within the early secretory pathway, vesicle coat proteins are key mediators of protein sorting, forming a cage-like protein structure around the transport vesicle [2]. Coat protein complex I (COPI) vesicles function in both, Golgi to ER retrieval and intra-Golgi trafficking, and thus maintain the correct localisation of ER- and Golgi-resident chaperones, enzymes and receptors. Anterograde (ER to Golgi) transport is operated by coat protein complex II (COPII) vesicles, budding from ER exit sites (ERES), distinct, ribosome-free ER regions [3]. The Sec23/Sec24 heterodimer is an early COPII component, recruited upon binding of the small GTPase Sar1 to the cytoplasmic face of the ERES [4]. Sec24 selectively binds to signal motifs of transmembrane cargo exposed to the cytoplasm. Luminal cargo, as soluble and membrane anchored protein, can be captured by Sec24 interacting transmembrane receptors at the ERES.

These general features of the secretory pathway are largely conserved in *Trypanosoma brucei*, a unicellular parasitic organism, causative of human and animal African trypanosomiasis and belonging to the early branching Discoba lineage. The *T. brucei* surface is dominated by glycosylphosphatidylinositol (GPI)-anchored proteins. In the bloodstream form (BSF), the lifecycle stage that persists in the intravascular system of the mammalian host, approximately 10^7^ variant surface glycoprotein (VSG) molecules per cell form a dense glycosylated surface coat [5]. The surface of the procyclic form (PCF), colonizing the *Glossina* spp. insect host midgut is radically different: The major surface protein are the procyclins, small polyanionic, GPI-anchored glycoproteins [6]. Given the substantial burden for folding and export of surface glycoproteins it is unsurprising that ER quality control (ERQC) pathways are present in *T. brucei*, including the luminal ER-associated degradation pathway (ERAD-L) that serves to target misfolded secretory proteins to degradation [7]. However, a canonical unfolded protein response (UPR) - the crucial adaptive stress signalling pathway of opisthokonts - is absent in trypanosomes. The canonical UPR is initiated by membrane-integral ER stress sensors including the protein kinases IRE1 and PERK and the transcription factor ATF6, each triggering separate signal transduction cascades that culminate in transcriptional reprogramming [8]. Trypanosomes rely on polycistronic transcription and lack transcriptional regulation of individual genes [9], and the absence of IRE1 and ATF6 homologs rules out a conventional UPR. Nevertheless, *T. brucei* encodes a PERK-like eIF2α kinase, PK3 (TbIF2K3), that phosphorylates eIF2α and induces the spliced leader RNA silencing (SLS) pathway in response to ER stress [10, 11]. Unlike the canonical UPR, which drives adaptive transcriptional reprogramming to restore ER homeostasis, SLS silences the spliced leader RNA required for trans-splicing of all mRNAs, leading to global translational shutdown; a terminal rather than adaptive response. How BiP activity is regulated to match secretory demand in trypanosomes, in the absence of canonical UPR sensors, therefore remains poorly understood. The core ERAD-L pathway and its effectors BiP, calreticulin, protein disulfide isomerases (PDIs) and mannose binding lectins, are well conserved in trypanosomes [12]. BiP belongs to the heat shock protein 70 (Hsp70) family and plays a key role in protein folding and ERQC. BiP carries the C-terminal ER-retention signal KDEL in mammals (MDDL in *T. brucei*). The KDEL receptor ERD2 specifically binds to luminal cargo molecules carrying such sorting signals in the ER–Golgi intermediate compartment (ERGIC) or cis-Golgi [13], facilitating retrograde transport to the ER. It has been demonstrated in *T. brucei* that removal of the C-terminal tetrapeptide allows BiP to escape the ER and enter the secretory pathway [14]. Further truncation of the entire C-terminal protein binding domain generates the secretory reporter BiPN which shows enhanced secretion [14].

To characterise the ER luminal proteome under the biosynthetic conditions of both, the mammalian-infective BSF stage, as well as the PCF stage, we exploit the differential trafficking of BiP and the truncated form, BiPN, as TurboID baits to derive a quantitative proximity proteome of the *T. brucei* secretory pathway that discriminates between ER, ER-to-Golgi transit and Golgi-resident components. Through strict ER retention, BiP labels other ER resident proteins and cargo proteins routed to the secretory pathway. Comparing the labelling from BiP and BiPN baits would result in a similar biotinylation pattern within the ER lumen but BiPN labelling activity additionally reveals Golgi proteins. The resulting data delivered a protein inventory of the secretory pathway, encompassing PCF and BSF life cycle stages, and revealed novel candidate ER, ER-to-Golgi transit and Golgi-resident proteins.

Additionally, we extended our approach to the nuclear membrane, which is continuous to the ER, while the inner nuclear membrane constitutes a specialised domain, distinct from the outer nuclear membrane. Employing the transmembrane nucleoporin NUP65, the inner ring nucleoporin that anchors the trypanosome nuclear pore complex to the nuclear envelope [15], as TurboID bait, we derived a protein repertoire of the inner nuclear membrane.

Lastly, we leveraged the BiP proximity-labelling derived ER proteome data to identify a new key player in trypanosome ER stress moderation, the SAP-domain containing protein Tb927.5.1160 for which we propose a function as BiP nucleotide exchange inhibitor (NEI) based on homology to the mammalian NEI MANF (mesencephalic astrocyte-derived neurotrophic factor) and its marked effect on the sensitivity towards ER-stress.

## RESULTS AND DISCUSSION

### A TurboID bait protein engineered to escape the ER

We chose the abundant ER chaperone BiP as bait for TurboID proximity labelling of the ER lumen. A sequence coding for a double hemagglutinin epitope-tag (2HA) followed by TurboID was genomically inserted into one allele of the BiP gene (Tb927.11.7510), upstream the coding sequence of the BiP C-terminal ER-retention signal-tetrapeptide (MDDL). Clonal lines from *in situ*-tagging (BiP-2HA-TurboID-MDDL, termed BiP-TurboID) were confirmed by western blotting (**Figure 1A and 1B**). Removal of the C-terminal MDDL signal leads to loss of ER-retention and secretion which could be elevated by the truncation of the entire C-terminal binding domain [14]. We selected this secretory reporter, termed BiPN, as a TurboID bait, fusing 2HA-TurboID to BiP residue 415 for tetracycline inducible expression (**Figure 1A)**. The 81 kDa BiPN fusion protein could be readily detected by western blotting (**Figure 1A and 1B**).

**Figure 1.**
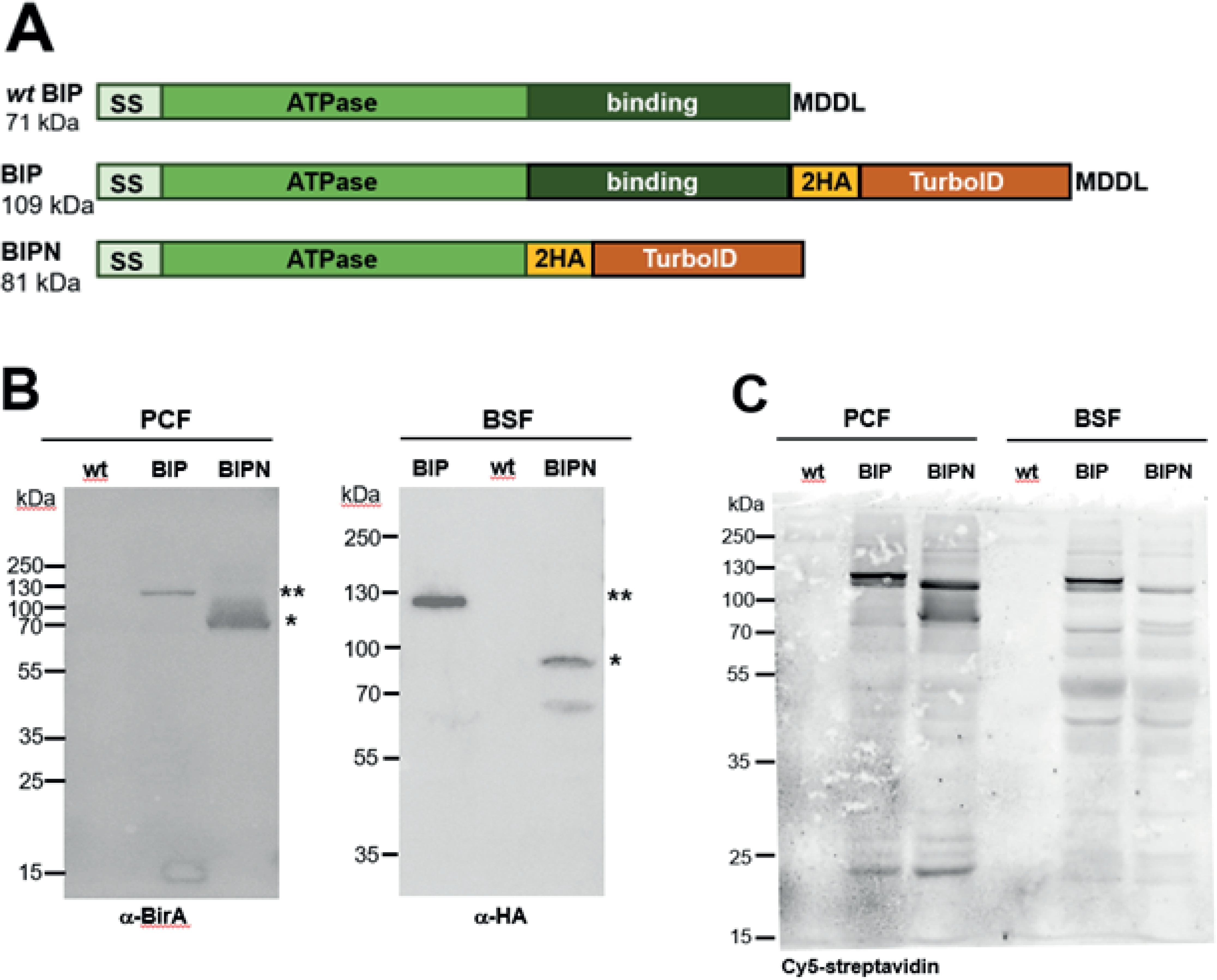
Generation of BiP and BiPN TurboID cell lines. (A) Schematic representation of domain and signal peptide organisation for the BiP and BiPN TurboID constructs as compared to wt BiP (SS = Sec signal; MDDL = ER retention signal; 2HA = duplicate HA epitope tag). (B) Western blot analyses of BiP and BiPN in BSF and PCF *T. brucei* using antibodies detecting the HA-epitope tag or the biotin ligase (BirA). The theoretical molecular weight of the fusion constructs is indicated by asterisks. (C) Cy5-streptavidin blot analysis of BiP and BiPN whole cell lysates in BSF and PCF *T. brucei*, respectively.

The BiP- and BiPN-TurboID lines were generated in both major life cycle stages, PCF and BSF trypanosomes. A streptavidin blot showed that all respective bait cell lines produced a distinct labelling pattern (**Figure 1C**). Among the most prominent bands are the respective bait proteins themselves. For BSF, a strong broad band running at approximately 50 kDa, likely originating from biotinylated VSG, is absent in PCF.

Next, we confirmed correct localisation of the two TurboID baits by fluorescence microscopy. The anti-BiP immunofluorescence signals in both TurboID strains are consistent with the wt strain in both life stages (**Figure 2 for PCF; Figure S1 for BSF**). The biotin modification from TurboID proximity labelling can be efficiently localized by fluorophore-labelled streptavidin [16]. Deployment of this probe on these specimens, produced a strong signal in all TurboID strains that largely colocalized with the anti BiP signal and is absent in parental cells (**Figure 2 for PCF; Figure S1 for BSF**). In PCF, we also tested fluorophore labelled ceramide which integrates preferentially into the ceramide-rich membranes of the TGN [17] and observed no significant colocalization with the streptavidin signal in this more distant Golgi compartment (**Figure S2A**). However, a co-stain of the streptavidin probe with antibodies detecting the Golgi-stack marker GRASP [18–20] showed that the biotinylation extends to the Golgi (**Figure 2B**). The bait proteins detected via HA-epitope also overlapped with the GRASP signal suggesting their presence at the ER exit sites (ERES) and the cis-Golgi (**Figure S2B**). In summary, we find a similar localization and labelling pattern for the BiP and BiPN which is expected as the BiP bait, despite being retained in the ER, shuttles between cis-Golgi and ER, and labels cargo proteins in the ER that are trafficked to the Golgi apparatus.

**Figure 2.**
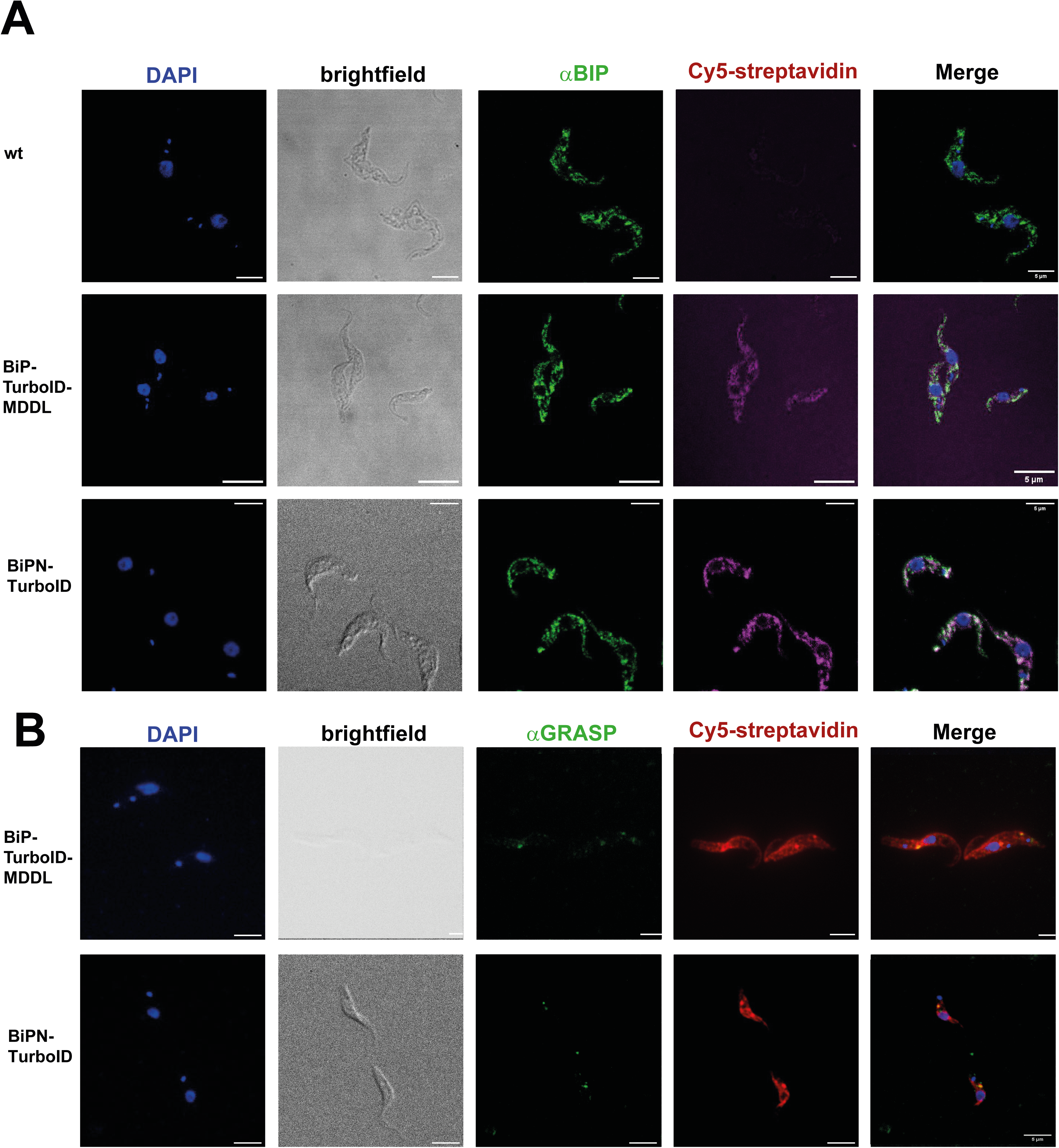
Localisation of TurboID fusion proteins in PCF T. brucei. (A) In *T. brucei* PCF, biotin labelling by BiP-TurboID fusion and BiPN-TurboID was detected with Cy5-streptavidin (red). The signal largely colocalises with that of anti-BiP antibody (green; Alexa488) consistent with an ER stain. Shown are single plane raw images for DAPI (blue), anti-BiP, Cy5-streptavidin fluorescence and a respective merge. Scale bar = 5 μm. (B) Biotinylation elicited by BiP- and BiPN-TurboID proximity labelling in PCF partly colocalizes with the Golgi apparatus in addition to the ER, as monitored by co-staining with Cy5-streptavidin (red) and anti-GRASP (green; Alexa488). Scale bar = 5 μm. Respective localisation of BiP-TurboID fusion and BiPN-TurboID in BSF are shown in **Figure S1**.

### BiP proximity labelling maps the luminal proteome of the *T. brucei* endoplasmic reticulum

The BiP and BiPN TurboID lines in the two life cycle stages were subjected to streptavidin affinity purification followed by LC-MSMS analysis, as previously described [21] and parental lines served as controls. For PCF and BSF 366 and 428 protein groups were found enriched in the respective BiP bait samples (**Figure 3A; Table S3**), 211 of which were shared. Based on TrypTag [22] and LOPIT [23]) annotation, the combined enriched cohort contained 106 ER, 16 Golgi, 68 endocytic, 46 surface and 5 acidocalcisome proteins (**Figure 3B; Table S3**).

**Figure 3.**
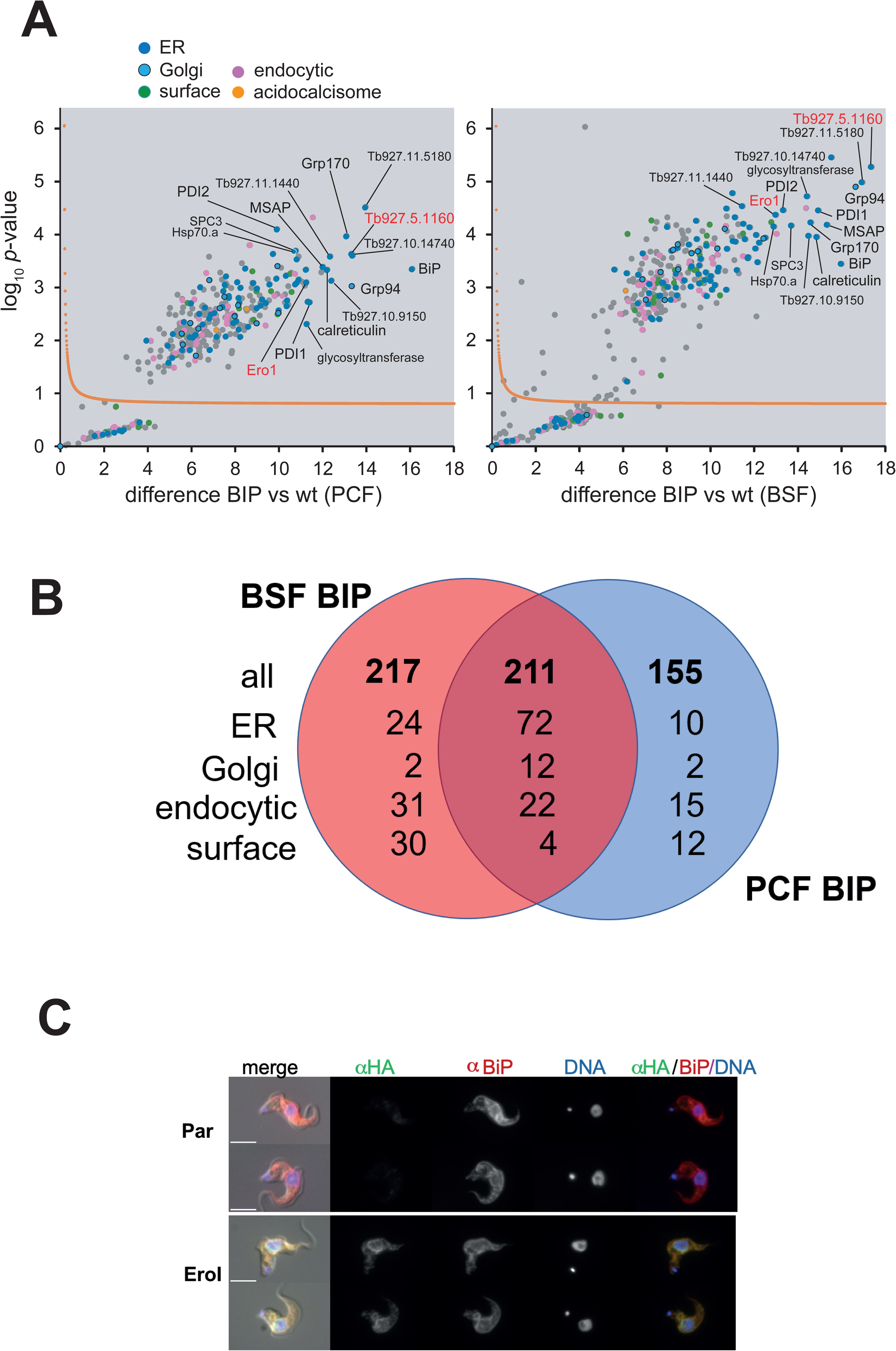
BiP TurboID. (A) Volcano plots for statistical analysis of BiP-TurboID experiments in PCF (left) and BSF *T. brucei* (right), generated by plotting the − log_10_*P*-value versus the *t*-test difference, comparing the BiP bait experiment to the parental control. A statistical cut-off curve (see Methods section) is drawn in orange. Protein groups assigned to ER (blue), endocytic (pink), surface or acidocalcisome based on TrypTag or LOPIT data [22, 23] are coloured. Selected proteins are labelled, the ER-localisation of proteins with red labels was validated by imaging in this work. SPC3, Tb927.5.1930; Grp94, Tb927.3.3580; Grp170, Tb927.9.9860; MSAP, Tb927.11.15210; Glycosyltransferase, Tb927.3.4630. (B) Venn diagram comparing the 583 BiP-TurboID enriched proteins for the two life-cycle stages. (C) Endogenous tagging confirms ER localisation of EroI (Tb927.8.4890) in BSF *T. brucei*. Anti-HA immunofluorescence of endogenously 6xHA-tagged EroI co-stained with anti-BiP antibody. Scale bars = 5 μm. For validation of tagging by western blot see **Figure S5**.

Overall, the detections robustly encompass prototypic, known ER proteins such as chaperones, the prototypic BiP interactor calreticulin (Tb927.8.7410) and PDIs (summarised in **Table S2**). Next to BiP auto-biotinylation, the strongest intensities, for both life stages, are found for the potential MANF (mesencephalic astrocyte-derived neurotrophic factor) homolog Tb927.5.1160 (see below), the Hsp90 chaperone Grp94 (Tb927.3.3580), the trypanosome Grp170 homolog (Tb927.9.9860) and five uncharacterised proteins. The latter appear to have trypanosome specific sequences, but structure predictions detected similarities to eukaryotic ER factors: Tb927.10.14740 carries a DnaJ2 J-domain motif (Prosite: PS50076; residues 224-294). Tb927.11.1440 shares similarities with the ER-resident thiol-disulfide oxidoreductase Sep15 from *Drosophila melanogaster* [24] (**Figure S3A, B**). The remaining three proteins, Tb927.11.5180, Tb927.10.9150 and Tb927.11.15210 (MSAP), resemble pERp1, an ER resident protein in mammals crucial for immunoglobulin and integrin folding [25]. All three appear to share the characteristic pERp1 disulfide bond pattern (**Figure S3C-E**) and have been previously localised to the ER [22] [26]. Lastly, the signal peptidase component SPC3 Tb927.5.1930 [27] is part of the strongest enriched cohort. In yeast the signal peptidase complex, facilitating cleavage of N-terminal signal sequences of proteins targeted to the secretory pathway, is composed of Spc1p, Spc2p, and Sec11p. The latter is the active signal peptidase Tb927.5.3220 [27], which is also labelled by BiP-TurboID.

In addition, we detected a plethora of known and uncharacterised proteins which have been localised to the ER by TrypTag or LOPIT [22, 23], as well as numerous surface proteins (summarised in **Table S4**) and secretory cargo targeted to organelles. A well characterized example of soluble ER cargo is the soluble lysosomal protease Cathepsin L (CatL). In trypanosomes, CatL delivery to the Golgi is p24 membrane receptor dependent [28]. Then, CatL is trafficked from the TGN to the lysosome, tied into clathrinlcoated vesicles. For this targeting step a dipeptide motif in the CatL prodomain has been identified, while the cognate signal receptor remains unknown [29]. Another example are proteins destined to the glycosome, peroxisome derived organelles with specialised metabolic functions in trypanosomes, that can form from vesicles budding from the ER [30]. BiP-TurboID enriches PEX11(Tb927.11.11520), a membrane bound peroxin and the related factor Gim5A that are involved in glycosome biogenesis [31]. Further BiP-TurboID detections are 31 protein groups with flagellar localisation, as for example the small GTPases Arl3A (Tb927.3.3450) and Arl3C (Tb927.6.3650) which have been implicated in ciliary functions in *T. brucei* [32], while the single mammalian Arl3 homolog localises to Golgi and cilium.

BiP-TurboID life-stage differences (**Figure S4, Table S3)** are most pronounced for surface proteins (**Table S4**) and largely consistent with life-stage proteomics analyses [33]. For the ER, a striking difference is for the ER degradation-associated mannosidase-related (EDEM) paralogs (Tb927.8.2940; Tb927.8.2930; Tb927.8.2920; Tb927.8.2910) that all exhibit stronger labelling in PCF. Mammalian BiP functions in the ERAD-L pathway together with PDIs, calnexin/calreticulin and interacts with the mannose-binding lectin Yos9 (Tb927.11.10700) [12], as well as with EDEM1 and ERdj5, a DnaJ variant with disulfide reductase activity carrying an N-terminal J domain and six thioredoxin domains at the C-terminus [34]. PDIs also display stage dependence, with PDI2 (Tb927.10.8230), previously reported as BSF-specific [35], showing higher levels (∼3 fold) in BSF, as does PDI1 (Tb927.4.2450) and Tb927.7.1300 (∼2 fold), while Tb927.5.1020 is increased in PCF (∼3 fold). The short PDI Tb927.7.5790, suggested a homolog of mammalian Erp44 [12] which competes with other PDIs for interaction with ER oxireductin (Ero1) in an ER stress dependent manner [36], is strongly enriched in BSF, again, consistent with life-stage proteomics data [33]

To independently confirm that BiP-TurboID identifies proteins with established ER function, we endogenously tagged EroI (Tb927.8.4890), a *T. brucei* ER oxidoreductin required for oxidative protein folding and VSG biosynthesis [12], with a 6xHA epitope at its C-terminus in BSF *T. brucei* by CRISPR-Cas9 using the pPOT7 donor system [37] (**Figure S5**). Anti-HA immunofluorescence revealed a reticulate signal consistent with ER morphology and extensive co-localisation with BiP (**Figure 3C)**, concordant with hyper LOPIT spatial proteomics which assigns EroI to the ER with probability 1.0 in both life cycle stages [23].

### TbMANF (Tb927.5.1160) is a constitutive ER component with a functional role in ER stress tolerance

Among the cohort of proteins that display the highest degree of labelling by BiPTurboID was Tb927.5.1160, a 26 kDa arginine-rich protein. BlastP searches and homology modelling suggest partial homology to MANF (**Figure 4**). Its mammalian counterpart is a known BiP interactor that binds to the ATPase domain via its SAP domain (spanning residues 128 to 169 in *M. musculus* MANF; compare **Figure 4A**) and inhibits BiP ADP exchange thereby stabilizing BiP client interactions, a constitutive property of MANF that attenuates BiP ATPase cycling independent of UPR activation [38]. The MANF region containing the SAP domain, including the structural elements engaging in BiP interaction, are conserved in Tb927.5.1160 (**Figure 4B)** which renders the protein a candidate BiP NEI. AlphaFold3 models of respective complexes with full-length BiP (including the BiP client-binding domain (CBD)) recapitulate the interaction surfaces between SAP domain and BiP ATPase domain that are relevant for NEI function (**Figure S6**). Notably, in these predictions the unrelated additional domains of MANF (a SAP-like domain) and Tb927.5.1160 (compare **Figure 4A)** are both contacting the BiP CBD albeit in a different manner (**Figure S6**).

**Figure 4.**
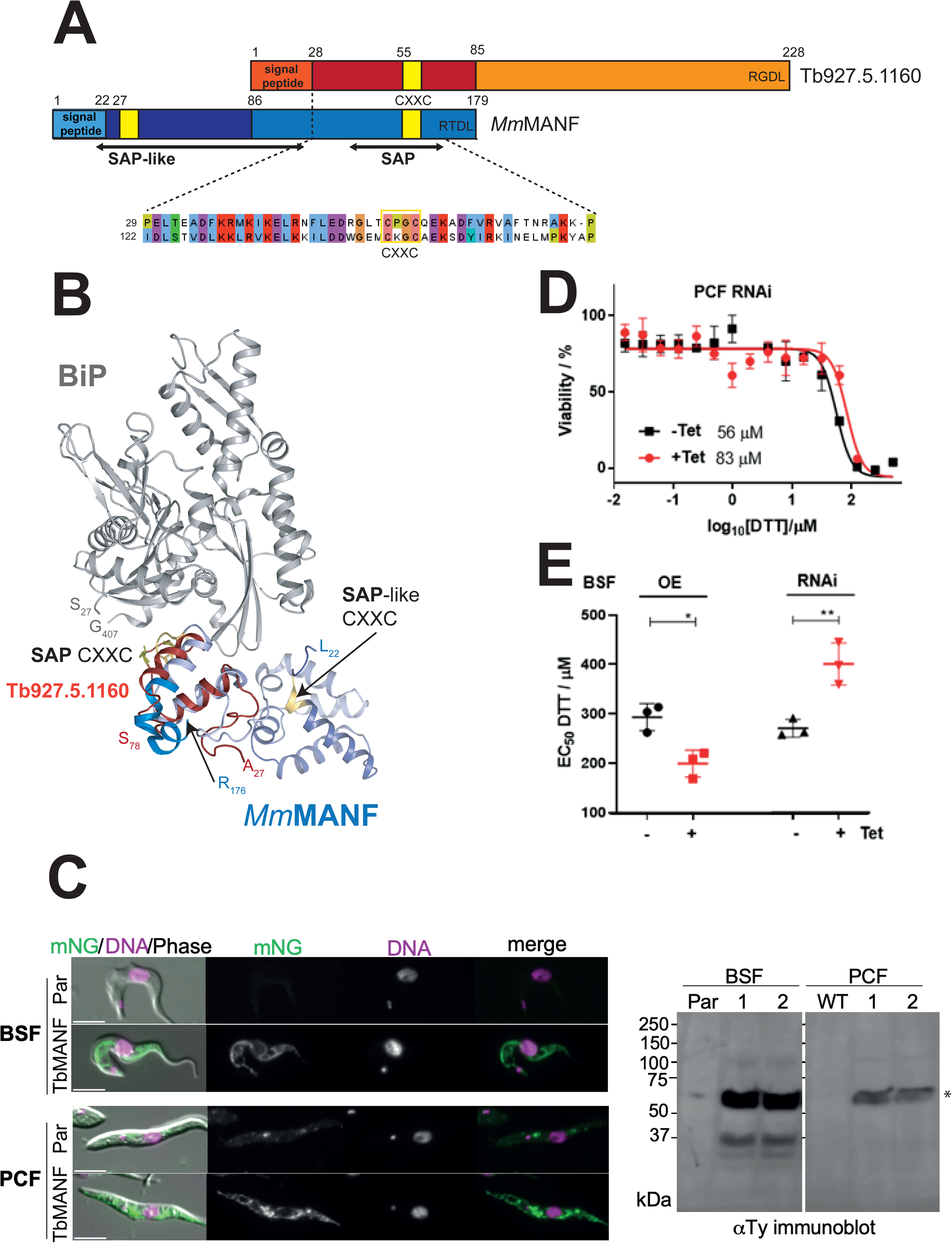
Tb927.5.1160 is a potential homolog of the mammalian BiP NEI MANF. (A) Schematic representation of domain and organisation of *Mus musculus* MANF and Tb927.5.1160. Saposin and saposin-like domains, both carrying a CXXC motif (yellow) and predicted signal peptides are indicated. (B) Cartoon representation of the crystal structure of the *Cricetulus griseus* BiP nucleotide binding domain (gray) in complex with murine MANF (blue) (pdb:6HA7) with the Phyre2 model of Tb927.5.1160 (red) superimposed. Comparison with a respective Alphafold3 model is shown in **Figure S6**. (C) Live-cell fluorescence microscopy of BSF and PCF *T. brucei* expressing endogenously tagged TbMANF::mNG::3xTy. Lightly fixed cells were imaged directly using the mNeonGreen fluorescence signal (green), co-stained with Hoechst 33342 to mark DNA (magenta). Parental (Par) and TbMANF-tagged cells are shown for each life cycle stage. Scale bars = 5 μm. Anti-Ty immunoblot (right) of BSF (parental, clones 1 and 2) and PCF (wild-type, clones 1 and 2) cell lines confirms tagged protein is detected at the expected molecular weight of approximately 66 kDa (theoretical molecular weight of indicated by asterisks). No band is present in parental cells. (D) RNAi depletion of Tb927.5.1160 in PCF *T. brucei* lowers the sensitivity towards DTT by approximately 1.5 -fold as determined by dose response analysis. EC_50_ values for the uninduced (-Tet) and induced RNAi (+ Tet) are indicated below the curves. (E) Tb927.5.1160 RNAi depletion in BSF *T. brucei* elicits a similar decrease in DTT sensitivity, while overexpression (OE) has the opposite effect. Shown are the average EC_50_-values determined from triplicate dose response analyses across three biological replicates with respective standard deviation. Unpaired *t*-test based significance is indicated by asterisks (* = *P*<0.1; ** = **P**<0.01). For detailed dose response data and growth curves see **Figure S7**.

In mammalian systems the nucleotide exchange factor Grp170 is stimulating ADP to ATP exchange on BiP [39] and murine MANF co-immunoprecipitates with BiP, Grp170 and the HSP90 family chaperon Grp94 [38]. Consistently, the *T. brucei* Grp170 and Grp94 homologs are highly enriched in BiP-TurboID along with Tb927.5.1160 (**Table S2**).

Tb927.5.1160 TrypTag annotation in PCF is ‘cytoplasm, reticulated’ [22] and its localisation in BSF is unknown. Given its strong enrichment in our ER proteome, we determined its localisation in both lifecycle stages without disrupting the endogenous 3’ UTR. We introduced a T2A-mNG::3xTy cassette at the endogenous Tb927.5.1160 locus in BSF and PCF *T. brucei* by CRISPR-Cas9 [40]. Correct integration was confirmed in two independent clones in each life stage by anti-Ty western blot, which detected a band at approximately 66 kDa absent from parental cells (**Figure 4C**). Live-cell fluorescence microscopy of lightly fixed BSF cells revealed a reticulate signal consistent with ER morphology (**Figure 4C**). In PCF we observed similar reticulated ER pattern, although Tb927.5.1160 abundance is substantially lower than in BSF [41]. The ER localisation is in agreement with LOPIT spatial proteomics, which assigns Tb927.5.1160 to the ER with probability 1.0 in both life cycle stages [23]. The endogenous tagging data confirm that Tb927.5.1160 is an ER-localised protein in both life cycle stages; consistent with its position as the dominant non-bait protein in the BiP proximal proteome.

Tb927.5.1160 RNAi knockdown in PCF led to a significant growth phenotype (**Figure S7A**), with whole cell proteomics reporting a target depletion to 13% upon 48h RNAi induction (**Figure S8A; Table S3**). Given that mammalian MANF constitutively modulates BiP ATPase cycling [38, 42], we tested the effect of the RNAi on ER-stress tolerance. Indeed, Tb927.5.1160 depletion led to an approximately 1.5-fold reduced sensitivity of PCF cells towards DTT (Figure 4D; **Figure S7A**). Next, we monitored DTT sensitivity in BSF *T. brucei* upon TbMANF depletion and respective inducible expression. Both, RNAi and inducible overexpression changed tolerance of ER-stress in opposite directions. While inducible expression led to significantly increased sensitivity towards dithiothreitol (DTT) (0.7 +/- 0.2-fold EC_50_, Figure 4D; **Figure S7C**), it is reduced in knockdown cells (1.4 +/- 0.1-fold EC_50_ change, Figure 4D; **Figure S7C**), similar to the PCF RNAi. This reciprocal response is consistent with TbMANF functioning as a modulator of BiP activity: elevated TbMANF stabilises BiP–client complexes by inhibiting ADP release, reducing the cell’s capacity to manage additional folding stress; conversely, reduced TbMANF permits faster BiP cycling, increasing folding throughput and buffering against DTT-induced disulfide disruption.

All three cell lines with manipulated TbMANF abundance were subjected to whole cell proteome analyses after 48h of induction using label-free, data-independent acquisition (DIA) quantitative proteomics. The BSF cell lines showed a 3.4-fold abundance increase of TbMANF for the induced expression, while RNAi depleted to 60% only (**Figure S8**). RNAi silencing in BSF led to the most severe growth phenotype (**Figure S7**) and the respective clonal lines were relatively unstable, which collectively could explain that only clones with mild depletion were obtained. Despite the marked growth phenotype of the depletion in PCF, the effect on the proteome was modest (**Figure S8A**): Along with the target, four other proteins were significantly decreased including a p24 transmembrane adaptor homolog (Tb927.11.5540) which detection is lost in the induced RNAi. A small 5.6 kDa uncharacterised Golgi protein (Tb927.11.14705) is among the three significantly increased proteins. Whilst there was similarly little proteome change upon Tb927.5.1160 overexpression in BSF (**Figure S8C, Table S3**), RNAi in this life cycle stage evoked a massive perturbation of the proteome landscape. Out of the 6600 proteins quantified almost half have significant abundance changes (2366 increased and 1224 decreased). Consequently, multiple GO terms are affected (**Table S3**), with predominant GO enrichment associated with ribosomes and ribosomal biogenesis in the upregulated cohort and those associated with mitochondrial and glycosomal metabolic pathways in the downregulated cohort. The ER GO component is enriched in the downregulated cohort albeit with a low enrichment factor of 1.6 (*P*<10^7^). These changes are likely not direct consequences of TbMANF depletion but strongly indicate secondary impact on cellular integrity. Altogether, the observed proteome changes fail to inform on TbMANF function but confirm essentiality in BSF *T. brucei*.

In conclusion, we describe here a novel ER factor in trypanosomes, acting proximal to BiP and exhibiting a conserved BiP binding region shared with the mammalian BiP NEI MANF. We further show ER-localisation in both life stages. While future *in vitro* studies are needed to confirm NEI function, we demonstrate that TbMANF is essential for *T. brucei* viability and has a significant role in the moderation of ER stress response. Mammalian MANF expression is stimulated in the course of the UPR [42]. Notably, TbMANF is the first candidate modulator of ER stress tolerance identified in trypanosomes, a lineage that lacks the canonical UPR sensors through which BiP activity is regulated in other eukaryotes.

### BiPN TurboID identifies known and prospective Golgi proteins with recognisable cytoplasmic sorting signals and secretory cargo

The BiPN construct lacks the ER retention signal and the C-terminal terminal substrate-binding domain, and is ultimately secreted, transiting from the ER through ERES and Golgi compartments before exiting the cell [14]. Proteins enriched in BiPN-TurboID relative to BiP-TurboID therefore should include factors encountered along this transit route: ERES-associated factors, Golgi-resident proteins, and secretory cargo co-trafficking with BiPN. Thus, BiPN-TurboID enrichment identifies proteins most likely to be Golgi-proximal but cannot by itself distinguish stable Golgi residents from transient trafficking factors and cargo. A direct comparison of BiPN with BiP TurboID (**Figure 5**) revealed the expected differential labelling with known experimentally localised Golgi proteins showing preferential BiPN labelling: In PCF, 13 out of 14 proteins with TrypTag Golgi localisation exhibit higher labelling in BiPN, including glycosyltransferases, a glycosyl hydrolase, adaptin-3 beta and the centrin-arm associated protein CAAP1. In BSF, this is the case for 9 out of 13 proteins. Additionally, the BiPN bait enriched the two highly divergent, Golgi resident N-acetylglucosaminyltransferases, GT11 (GnTI) and GT15 [3, 43, 44], in BSF, the stage these are predominantly expressed [33, 45]. The detection level in PCF is too low for a reliable quantification. Tb927.3.5330, a homolog of the GTPase activating protein centaurin β5, localized to the Golgi and nucleolus by Tryptag (nuclear localization by LOPIT), also shows no BiPN TurboID enrichment, but the nuclear localisation may indicate that the protein is cargo rather than Golgi resident.

**Figure 5.**
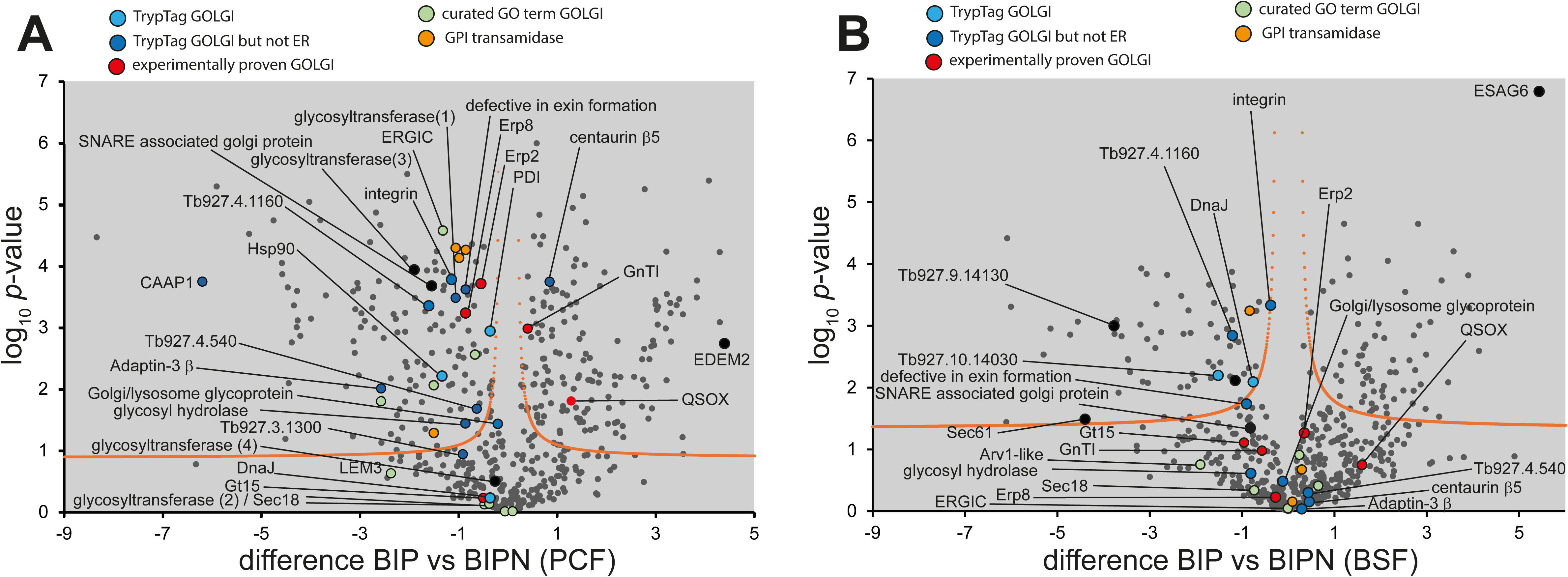
BiPN-TurboID enriches proteins encountered during ER-to-Golgi transit, including Golgi-resident proteins. (A) Volcano plots for comparison of the BiP-TurboID (right side of plot) and BiPN-TurboID (left side of plot) experiments in PCF (A) and BSF (B) *T. brucei* (right), generated by plotting the − log_10_*P*-value versus the *t*-test difference, comparing the two baits. Protein groups with TrypTag Golgi localization are in blue (dark blue for ER and Golgi; light blue for Golgi) and those experimentally validated are in red. Additional proteins with curated GO component annotation ‘Golgi Apparatus’ are in green, GPI transamidases are in orange and further proteins discussed in the text are in black. A statistical cut-off curve is drawn in orange. For complete annotation see **Table S3.** Golgi/lysosome glycoprotein, Tb927.8.1870; glycosyl hydrolase, Tb927.11.10370; ERGIC, Tb927.11.4200; glycosyltransferase (1), Tb927.10.12290; glycosyltransferase (2), Tb927.4.5240; glycosyltransferase (3), Tb927.6.1140; glycosyltransferase (4), Tb927.2.2380; GnTi, Tb927.3.5660; Gt15, Tb927.7.300; DnaJ, Tb927.11.16740; Hsp90 (Grp94), Tb927.3.3580; LEM3, Tb927.9.12000; QSOX (quiescin sulfhydryl oxidase), Tb927.6.1850; SEC61, Tb927.11.6230; Sec18, Tb927.11.1680; SNARE associated golgi protein, Tb927.11.8060; adaptin-3 beta, Tb927.11.10650; Centrin arm-associated protein CAAP1, Tb927.10.1450

Protein components of ERES, including cargo receptors, as for example the p24 transmembrane adaptor homologues Erp2 (Tb927.9.15090) and Erp8 (Tb927.10.6640) [28] also showed increased labelling by BiPN in PCF (**Table 1**), consistent with these factors concentrating at the ERES, the first compartment accessed by the secreted BiPN reporter. The same is true for the putative ER–Golgi intermediate compartment (ERGIC). ERGIC53 (Tb927.11.4200), a transmembrane lectin which binds glycoprotein cargoes and facilitates their loading into COPII-coated vesicles through interaction at the cytoplasmic side in mammals [46] thereby exerting selective quality control [12]. ERES are in close proximity of the flagellar attachment zone (FAZ)-associated ER [3] which may explain the detection of several FAZ components, as for example FLA1 and FLA1-binding protein, protein components of the FAZ which bridge the connection between flagellar and cytoplasmic membrane [47] and a VAMP-associated protein (Tb927.11.13230) which contributes to FAZ-associated ER maintenance [48]

**Table 1.**
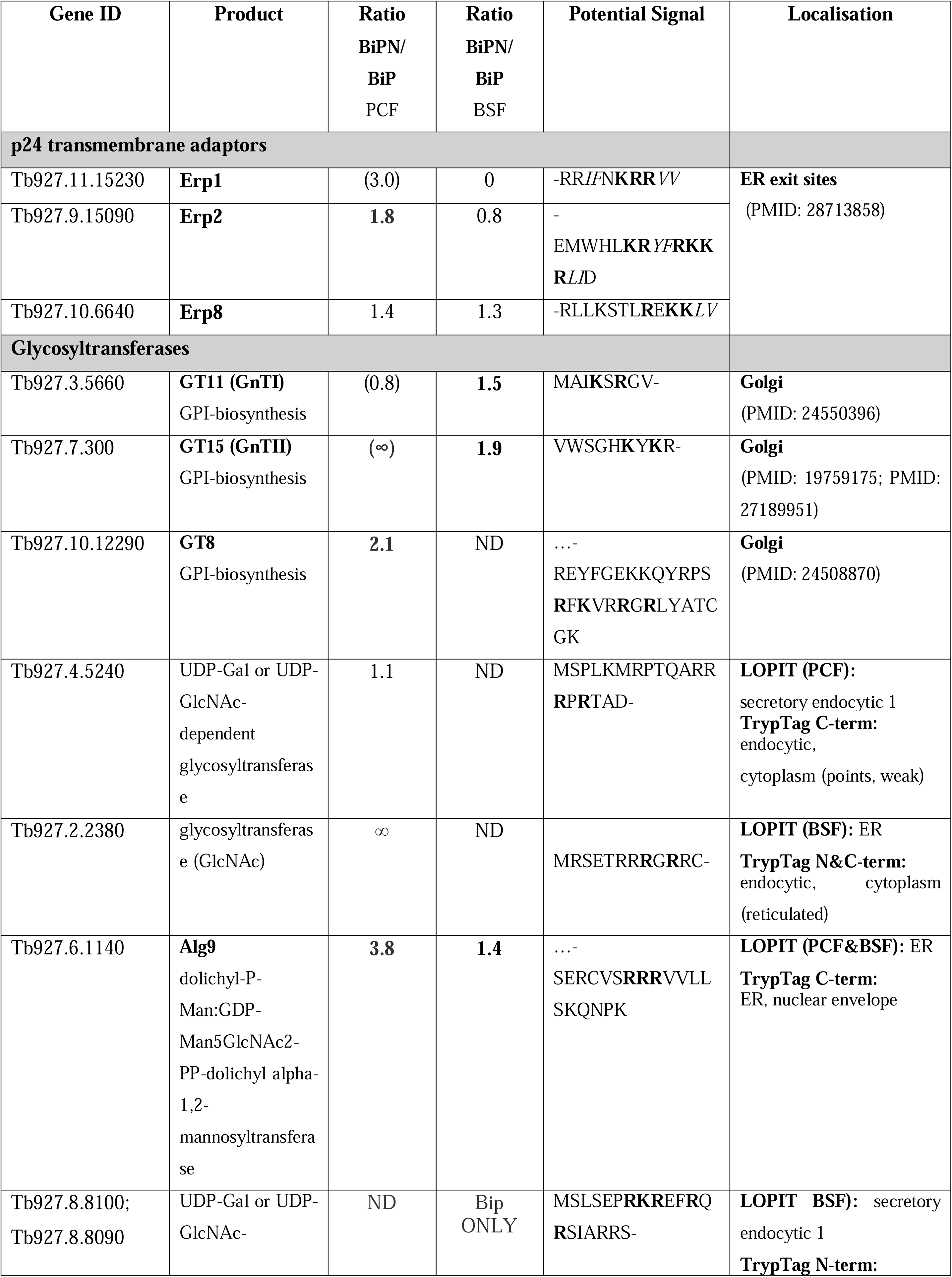
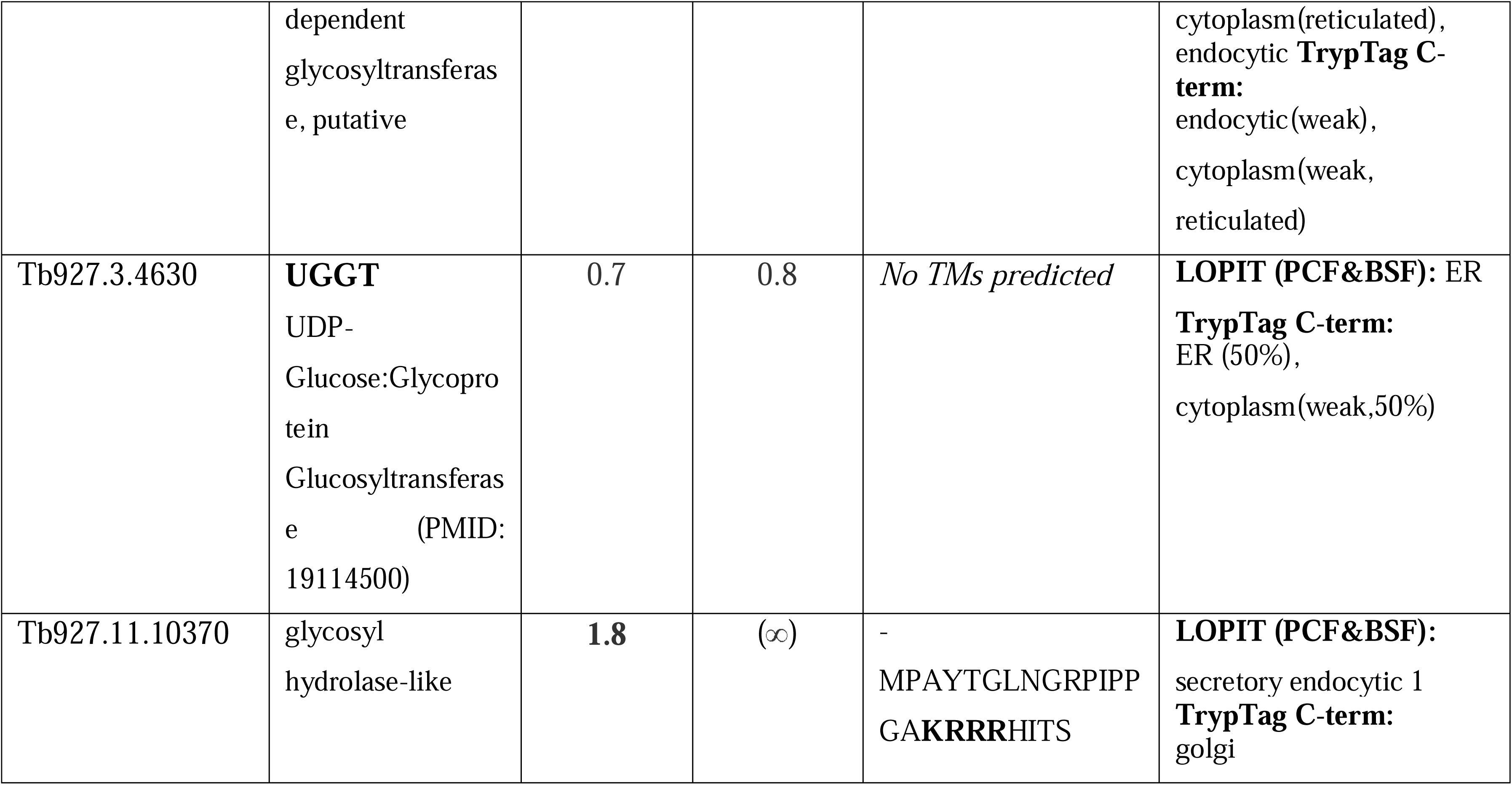
Summary of p24 transmembrane adaptors and glycosyltransferases detected [80]. Ratios from LFQ intensities of BiPN and BiP are given for both life cycle stages and potential COPI/COPII signals at the predicted cytosol exposed terminus are shown. For p24 transmembrane adaptors such signals were previously proposed[28]. Ratio values in brackets are based on borderline detections and are thus low confidence.

Whilst the above comparison with known Golgi factors offers proof of concept for our BiPN TurboID approach, we also observed a certain bias: Firstly, as expected for a transit-based reporter, labelling efficiency appears to vary across Golgi sub-compartments. An example, common between the two life stages, is quiescin sulfhydryl oxidase (QSOX), which resides in the Golgi [11]. QSOX shows weaker BiPN enrichment, likely reflecting its concentration in distal Golgi cisternae where the local density of transiting BiPN may be lower.

Secondly, a potential weakness of the BiPN TurboID approach is that BiP client proteins which interact with BiP in the ER exhibit decreased labelling because of the absent CBD in BiPN. This is exemplified by DnaJ family members, known to interact with mammalian BiP while playing their role in targeting clients to BiP and modulating BiP’s ATP-dependent activity [49]. This ER resident DnaJ cohort indeed shows significantly stronger labelling with BiP (**Table S2**). A consistent exception is Tb927.11.16740, a DnaJ homolog with demonstrated Golgi and ER localisation [22]. Similarly, a cohort of heat-shock proteins (**Table S2**), including Grp94, for which both, direct interaction with BiP [50] and Golgi-presence [22] have been demonstrated, display higher enrichment in BiPN-TurboID. Moreover, the missing CBD should affect the differential BiPN/BiP labelling of client proteins, which would explain the observed variance of BiPN/BiP ratios between paralog groups of surface protein cargo (compare **TableS4**). For example, GP63 family surface protease members and trans-sialidases are preferentially labelled by BiP while invariant surface glycoproteins ISG75 and ISG65 exhibit stronger BiPN labelling, which may indicate lesser reliance on BiP chaperone function.

Altogether, these complexities preclude confident prediction of Golgi residency based on differential BiPN/BiP labelling alone. However, these data offer orthogonal support when used in conjunction with localisation data from other approaches and the presence of relevant COPI/COPII sorting signals. COPII mediated vesicle formation is a universal mechanism of ER exit in eukaryotes. In *T. brucei* ERES are situated proximal to the FAZ-associated ER [3]. The Sec23/Sec24 heterodimer is an early COPII component, recruited upon binding of the small GTPase Sar1 to the cytoplasmic face of the ERES [4]. Sec24 selectively binds to signal motifs of transmembrane cargo exposed to the cytoplasm. Luminal cargo, such as soluble and membrane anchored protein, can be captured by Sec24 interacting transmembrane receptors at the ERES. *T. brucei* encodes two sets of Sec23/Sec24 paralogs [51] and for the ERES p24 transmembrane adaptors a COPI/COPII signal, localised at the cytosol exposed C-terminus was proposed [28]. Here, [RK](X)[RK]XX serves for Golgi directed (anterograde) transport while a set of two hydrophobic patches facilitate interaction with COPI (**Table 1**). For Golgi resident glycosyltransferases, a dibasic motif [RK](X)[RK] located proximal to the transmembrane border has been described for COPII vesicle mediated transport to the Golgi [52]. This signal is detectable in the N-terminal cytoplasmic peptides of the N-acetylglucosaminyltransferases GnTI (MAI**K**S**R**GV-) and Gt15 (VWSGH**K**Y**K**R-) as well as in further glycosyltransferases that exhibit stronger BiPN labelling (**Table 1)**.

Overall, our BiPN/BiP proximity labelling approach delivers orthogonal evidence for protein localisation in sub-compartments of the *T. brucei* secretory pathway, complementing existing resources based on spatial proteomics and imaging.

### NUP65C-TurboID: The proteome of the inner nuclear membrane

In *T. brucei*, the protein composition of the inner nuclear membrane has not been systematically determined. The inner nuclear membrane is a specialised ER-continuous domain, that is distinct from the bulk ER and separated from the outer nuclear membrane by the nuclear pore complex. It has distinct functions in nuclear organisation and gene expression, and it is physically separated from the remaining ER by the nuclear pores. Proteins of the inner nuclear membrane are synthesised at the ER and subsequently transported to the inner NE, usually via the small peripheral channels of the pore close to the high-curvature-membrane [53, 54]. In general, this passage prevents inner NE proteins from having too large cytoplasmic/nuclear domains, and the high-curvature-membrane possibly also adds some further sterical restrictions. Apart from this, there is no positive or negative sorting to the inner NE at the pores, and many ER proteins constantly cycle through the inner NE [55]. The enrichment of specific proteins at the inner NE is instead mostly achieved by these proteins having high affinities to nuclear proteins or chromatin [53, 56–58](diffusion-retention mechanism, and by selective removal of the other proteins by the ubiquitin-ligase based INM-associated degradation (INMAD) [53]. In *T. brucei*, the composition of the inner nuclear membrane is unknown.

To specifically identify the proteome of the inner nuclear membrane, we extended our proximity labelling approach to this membrane domain and expressed *T. brucei* NUP65 with a C-terminal TurboID fusion. NUP65 is the inner ring nucleoporin that anchors the trypanosome nuclear pore complex to the nuclear envelope via a TM domain. It has a large N-terminal intracellular domain, facing the nuclear pore, followed by the TM domain and a C-terminal domain that reaches into the lumen of the nuclear envelope (**Figure 6A**). We reasoned that NUP65-TurboID will mostly biotinylate membrane proteins on their passage to the inner nuclear envelope, including both specific inner nuclear membrane proteins as well as ER proteins that pass non-specifically due to the absence of any negative selection. We labelled the cells with fluorescent streptavidin to visualise the biotinylation of NUP65-TurboID. The biotin signal was mostly at the pores and at the nuclear envelope, and much weaker at the endosomal system, confirming the correct integration of the fusion protein to the membrane (**Figure 6B**). In contrast, the biotinylation pattern of an N-terminal TurboID fusion was concentrated at the pores (**Figure 6C**).

**Figure 6.**
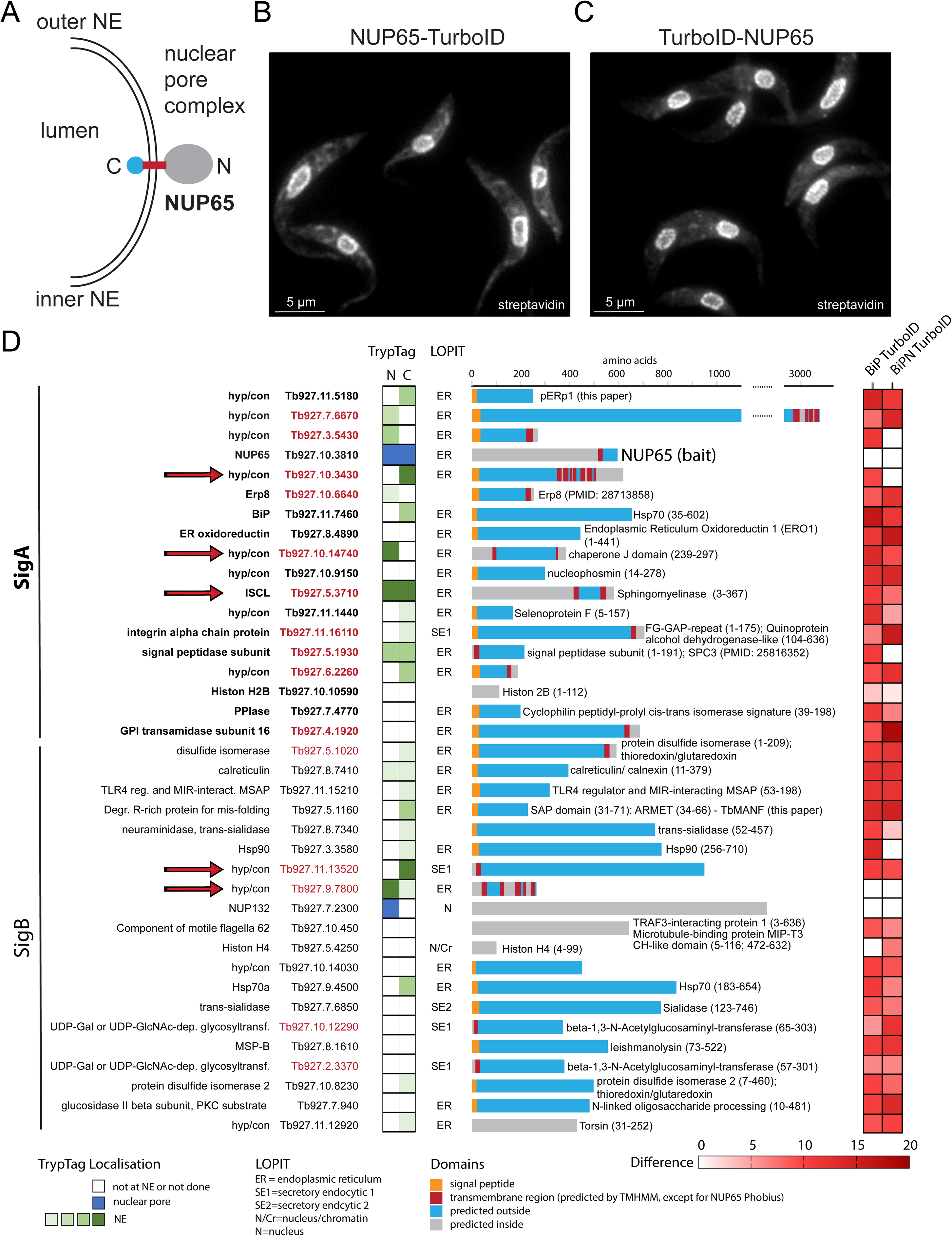
A proteome of the inner nuclear membrane. Localisation (schematic) of the inner ring NUP65 to the nuclear membrane (A). Cells expressing NUP65 fused to TurboID either at the C-terminus (B) or N-terminus (C) were labelled with fluorescent streptavidin. Single plane, representative images are shown as raw data. List of all proteins (SigA and SigB) labelled by NUP65-TurboID (D). The localisation to the NE has been manually evaluated looking at the raw images of C or N terminal tagged GFP fusions from TrypTag [22] and classified in different shades of green, according to the strength and dominance of the NE localisation. LOPIT data are presented [23] as well as schematics of the proteins with their predicted signal peptides, transmembrane helices, intra-and extracellular domains (DeepTMHMM-1.0, [78]. Additional domains are shown [79]. A heatmap (right) indicates the degree of eaznrichment observed for BiP- and BiPN-TurboID in PCF for each protein (the shade of red corresponds to the statistical *t*-test difference as compared to a wt-control).

Next, we purified the biotinylated proteins by streptavidin affinity and analysed the peptides by mass spectrometry. We defined 18 proteins as highly significantly enriched (SigA, FDR = 0.01, s0 = 0.1; this included the bait NUP65) and 20 as significantly enriched (SigB, FDR = 0.05, s0 = 0.1), in comparison to WT cells (**Figure 6D**). Of the SigA proteins, all proteins except NUP65 were detected by BiP-TurboID and 15/18 proteins have ER localisation predicted by LOPIT [23], one is secretory endocytic. Of the two remaining proteins without LOPIT data in PCF, Erp8 was experimentally localised to the ERES [28]. Only one protein, histone H2B, cannot be possibly accessed by NUP65 and appears to be a contaminant. Excluding the bait, ten of these proteins have at least one predicted TM domain and are thus possible proteins of the inner nuclear membrane (red font in **Figure 6D**). In fact, all have a rather small (predicted) intracellular domain, consistent with a potential negative selection of bulky proteins at the peripherical pore channels. To evaluate whether these 10 proteins are specific components of the inner NE or just ER proteins with no negative sorting at the pore, we explored the genome wide localisation database TrypTag [22]. With the exception of one protein (an essential GPI transamidase subunit 16), all proteins appear to be localised to the nuclear envelope to some extent. For three proteins, NE localisation was very dominant, indicating a positive selection (red arrows in **Figure 6D**). Tb927.10.3430 is a hypothetical protein with seven transmembrane helices, but no other recognisable conserved domains. Tb927.10.14740 is a hypothetical protein with 2 TM helices, flanking a large extracellular domain and a predicted chaperone J domain. Tb927.5.3710 is annotated as inositol phosphosphingolipid phospholipase C [59]. The proteins classified as SigB are also mostly proteins of the ER or secretory system according to LOPIT fractionation [23], with four exceptions that are likely contaminants. 3 out of these 20 proteins were not detected by BiP-TurboID, i.e., Histon H4, the nucleoporin NUP132 and Tb927.9.7800. Only five proteins contain TM domains, and of these, two proteins have a dominant localisation to the NE, the hypothetical proteins Tb927.11.13520 (one TM) and Tb927.9.7800 (5 TMs). Neither of these two proteins has any recognisable additional domain.

In summary, our NUP65-TurboID approach has identified almost exclusively proteins of the ER or endocytic system, with only 4 putative contaminants among 38 proteins and thus has a high specificity. About half of these 34 proteins with endocytic localisation had a predicted TM domain. Of these, 5 showed a significant enrichment at the NE in the Tryptag localisation project, indicating a positive selection. With one exception, all positively-selected proteins are hypothetical proteins with no conserved domains outside *T. brucei,* suggesting trypanosome-specific function that await characterisation.

## Conclusion

The *T. brucei* secretory pathway operates under extreme biosynthetic pressure and does so through mechanisms that diverge from mammalian models. Canonical UPR signalling effectors and ER quality control machinery remain largely uncharacterised, yet these parasites rely on delivery of GPI-anchored surface proteins. Expression of our TurboID proximity biotinylation bait protein in the ER lumen provides a route to characterising this machinery that is complementary to existing spatial proteomics approaches and well-suited to the discovery of ER-resident proteins regardless of their retention mechanism. The BiP and BiPN datasets presented here define the first comprehensive proximity proteome of the *T. brucei* ER lumen across both major life cycle stages, capturing established ER quality control factors, secretory cargo, and novel candidate Golgi proteins whose Golgi targeting is supported by recognisable cytoplasmic sorting signals.

Amongst the proteins identified, TbMANF (Tb927.5.1160) is the most strongly labelled BiP-proximal non-bait protein and provides a first candidate regulator of BiP ATPase cycling in kinetoplastids. Because *T. brucei* lacks IRE1/ATF6/PERK, the three canonical UPR sensors that rely on regulated BiP interaction in mammals, TbMANF may represent an alternative, constitutive mechanism for regulating BiP activity in proportion to secretory demand. The essentiality of TbMANF across life cycle stages, and its higher abundance in the biosynthetically active bloodstream form, are consistent with this model.

Finally, the NUP65-TurboID inner nuclear membrane proteome demonstrates that the proximity labelling strategy is extendable to specialised ER-continuous compartments, identifying five candidate inner nuclear membrane proteins. These proteins are largely trypanosome-specific and represent an uncharted aspect of kinetoplastid nuclear organisation. Together, these datasets provide a foundation for systematic dissection of ER function, Golgi organisation, and nuclear envelope biology in an early-diverging eukaryotic pathogen.

## Methods

### Cell culture

Procyclic form (PCF) *Trypanosoma brucei brucei* (strain Lister 427) cells were cultured in SDM-79 media containing 10% heat-inactivated fetal bovine serum (FBS) at 27°C, maintaining a cell density below 1×10^7 cells/ml. Bloodstream form (BSF) cells were cultured in HMI-9 media containing 10% heat-inactivated FBS at 37°C in 5% CO_2_, maintaining a cell density below 1×10^6 cells/ml.

### Cloning and transfection

BiP was endogenously tagged to generate BiP-2HA-TurboID-MDDL utilising the PCR tagging based on pPOT7 [37]. pPOT7 was modified for TurboID-2xHA-tag fusions. BiPN-TurboID was expressed from the tetracycline-inducible plasmid p3383 [60] and the respective BiP fragment (to residue 415) was amplified from genomic DNA. Tetracycline inducible RNAi cell lines in *T. brucei* BSF and PCF, were generated using the pRPaiSL [61] or the p3666 [62] based stem-loop RNAi system, respectively, using primers predicted by RNAit2 [63]. Tb927.5.1160 inducible expression in *T. brucei* BSF relied on the plasmid construct pRPa^Tag^ and its integration into the tetracycline-responsive rRNA promoter of 2T1 cells [61] RNAi lines were confirmed by quantitative RT-PCR using the Maxima H Minus cDNA Synthesis Master Mix (Thermo) and Maxima SYBR green qPCR master mix (Thermo) and a Biorad CFX96 cycler with 95°C/10 min (initial denaturation), then 95°C/15 s (denaturation), 60°C/30 s (annealing), and 72°C/40 s (extension) over 40 cycles. *Tb*GAPDH served as endogenous reference for normalization. All primers are listed in **Table S1**.

### Endogenous protein tagging by CRISPR-Cas9

EroI (Tb927.8.4890) was tagged at its endogenous C-terminus with a 6xHA epitope in BSF *T. brucei* using the CRISPR-Cas9-based pPOT7 donor plasmids [37] with primers (Table S1) and G00 (sgRNA) scaffold generated as described in [64]. Correct integration was confirmed by anti-HA western blot prior to immunofluorescence. For co-localisation analysis, EroI-6xHA cells were fixed with 4% formaldehyde, permeabilised with 0.5% NP-40, and co-stained with rat anti-HA (clone 3F10; 1:1,000; Roche) and rabbit anti-BiP (gift from J. Bangs, University at Buffalo; 1:200), detected with Alexa Fluor 488-conjugated anti-rat and Alexa Fluor 594-conjugated anti-rabbit secondary antibodies respectively. Images were acquired on a motorised Zeiss AxioImager M2 stand (×100 PlanApo oil immersion objective, NA 1.46) equipped with a triple-pass DAPI/FITC/Texas Red emission cube using a Hamamatsu ORCA-Flash4 camera, and analysed in ImageJ v2.16.0/1.54p.

Endogenous C-terminal tagging of TbMANF (Tb927.5.1160) was performed in BSF *T. brucei* using the pRExT2A CRISPR/Cas9-based tagging system, which retains the endogenous 3′ UTR [40]. Briefly, donor repair templates were PCR-amplified from the pRExT2A-CT-HmNG plasmid (configuration: mNG::3xTy::T2A::HYG::3xTy) using gene-specific primers containing 30 bp homology arms targeting sequences immediately upstream of the stop codon and the adjacent 3′ UTR (**Table S1)**. For TbMANF, the repair cassette encoded mNG::3xTy linked via the Thosea asigna virus 2A (T2A) peptide to a hygromycin resistance marker for positive selection. Synonymous mutations were introduced into the repair template primers to disrupt the PAM site and prevent further Cas9 editing following integration, and primer design ensured that the full-length endogenous 3′ UTR remained intact downstream of the cassette insertion. Single guide RNAs (sgRNAs) were generated by PCR as described [40, 64], using gene-specific primers encoding a T7 promoter and 20 bp targeting sequence adjacent to the PAM site near the stop codon (**Table S1**). Pooled sgRNA and repair template PCR products were co-transfected into BSF *T. brucei* constitutively expressing SpCas9, and correctly tagged clones were selected with 5 μg ml^-1^ hygromycin and isolated by limiting dilution. Correct integration was confirmed by anti-Ty western blot, which detected a band at the expected molecular weight in three independent clones, that was absent from parental cells. Subcellular localisation of TbMANF::mNG::3xTy was assessed by live-cell fluorescence microscopy of lightly fixed cells as described [37].

### Immunofluorescence microscopy

PCF and BSF cells were harvested and washed twice with PBS, respectively. Cells were fixed with 4% paraformaldehyde on ice for 15 min and processed as described [65]. The TurboID-HA tag was detected with anti-BirA (rabbit polyclonal; PA5-80250; Invitrogen; 1:200 dilution) in combination with anti-rabbit IgG Alexa Fluor 488 (A-11008; Themo Fisher Scientific; 1:1000 dilution). Further antibodies used were rabbit anti-BiP (from J. Bangs, University at Buffalo; 1:200 dilution). Anti-rabbit GRASP (from G. Warren; 1:400 dilution) served as Golgi marker. Biotinylation was detected using Cy5- or Cy3- labelled streptavidin (Cytiva; 1:200 dilution), essentially as reported [16]. Images were acquired by using Leica a TCS SP8 WLL SMD-FLIM confocal microscope with a HC PL APO CS2 objective (63x Oil, NA 1.4, WD 0.14 mm, DIC) using Type F Immersion oil (NA:1.51) and LAS-X software (Leica Microsystems). The excitation and emission light was separated by filter sets NF488 (excitation 495nm–550nm) for Alexa Fluor™ 488 combined with SMD2 NF 405/640 (excitation 645-680) for Cy5-Streptavidin. Regarding with the Golgi complex staining, setting was used 405/488/532/647, Cy3 with 532 laser excitation and 566/590 emission commitment with 593/40 filter set. Images were recorded as Z-stacks of 30 to 50 images in 300 nm distances. Images were processed and analysed without deconvolution using Fiji software.

### Golgi lipids staining

The Golgi apparatus (TGN and trans-Golgi cisternae) was stained by BODIPY™ TR ceramide (D7540, Thermo Fisher Scientific), following the manufacturer’s instructions. Briefly, 1×10^7 PCF cells were pelleted, then resuspended in serum- and hemin-free SDM79 medium. Cells were treated with a final concentration of 5 µM of BODIPY™ TR ceramide for 1 h at 4°C, then washed thrice with ice-cold medium and incubated with fresh medium at 27°C for a further 30 min before proceeding with fixation, permeabilization, blocking and immunostaining.

### Western and streptavidin blotting

2×10^7 cells were harvested and washed once with PBS, before resuspending the cell pellet in 100 μl of SDS sample buffer containing 1 mM DTT. About 5×10^6 cells were loaded onto a 12% SDS-PAGE gel. Dilution of antibodies was as follows: 1:2000 dilution of rabbit anti-HA; 1:2000 dilution of rabbit anti-BirA antibody; 1:2000 dilution of Cy5-streptavidin; 1:15,000 dilution of Goat-anti-Rabbit IgG H+L HRP conjugate (AP307P, Sigma).

### Affinity purification of biotinylated protein, mass spectrometry and analysis

Biotinylated proteins were enriched by streptavidin Dynabeads (MyOne Streptavidin C1; Thermo Fisher Scientific) for 1 h at 4 °C under gentle mixing. Trypsin digestion and peptide preparation were performed as described [66] except that 1 mM biotin was added to the bead tryptic digests. Peptides were injected to reverse C18 column coupled to tandem mass spectrometry (LC-MSMS) with an Orbitrap Fusion mass spectrometer (Thermo Fisher Scientific). Spectra were processed and data normalized by using the intensity-based label-free quantification (LFQ) in MaxQuant version 2.0.9.0 [67, 68] searching the *T. brucei brucei* 927 annotated protein database (release 68) from TriTrypDB [69]. Data analysis was done using Perseus [70, 71]. LFQ intensities were log2-transformed and missing values imputed from a normal distribution of intensities around the detection limit of the mass spectrometer. A Student’s t-test was used to compare the LFQ intensity values between the triplicate samples of the bait with untagged control (WT parental cells) triplicate samples. The -log10 p-values were plotted versus the t-test difference to generate multiple volcano plots (Hawaii plots). Potential interactors were classified according to their position in the Hawaii plot, applying cut-off curves for significant class A (SigA; FDR = 0.01, s0 = 0.1) and significant class B (SigB; FDR = 0.05, s0 = 0.1). The cut-off is based on the false discovery rate (FDR) and the artificial factor s0, which controls the relative importance of the t-test p-value and difference between means (at s0 = 0 only the *P*-value matters, while at non-zero s0 the difference of means contributes).

### Whole cell proteomics

PCF or BSF cells were induced with tetracyclin (1 μg/ml) for Tb927.5.1160 RNAi depletion or inducible expression and incubated for 48 h in parallel with nontreated cells. 5 x 10^7^ cells were harvested within their logarithmic growth phase and resuspended in PBS containing Complete Mini Protease Inhibitor Mixture (Roche), then washed twice in PBS. Cells were then mixed with 100 mM TEAB (Triethylammonium bicarbonate) containing 2% SDC (sodium deoxycholate) and lysed by boiling at 95°C for 5 min and sonication (Bandelin Sonoplus Mini 20, MS 1.5). Protein concentration was determined using BCA protein assay kit (Sigma Aldrich). 20 µg of protein was adjusted with 100 mM TEAB (Triethylammonium bicarbonate) containing 2% SDC (sodium deoxycholate) to a final concentration of 50 µl, mixed with 40 mM chloroacetamide, 10 mM TCEP (Tris(2-carboxyethyl) phosphine) and boiled again. Samples were then mixed with 5 µl of SP3 beads (Hughes et al., 2019) and 65 µl of 100 % ethanol. After binding, the beads were washed twice with 80% ethanol. The mixture was digested in 100 mM TEAB with 0.5 µg of trypsin overnight at 37°C. After digestion, samples were acidified with TFA to 1% final concentration and peptides were desalted using in-house made stage tips packed with C18 disks (Empore).

Nano Reversed phase columns (Ion Opticks, Aurora Ultimate TS 25×75 C18 UHPLC column) were used for LC/MS analysis. LC was performed on a Vanquish NEO UHPLC system (Thermo Scientific). Mobile phase buffer A was composed of water and 0.1% formic acid. Mobile phase B was composed of acetonitrile and 0.1% formic acid. Samples were loaded onto the trap column (C18 PepMap100, 5 μm particle size, 300 μm x 5 mm, Thermo Scientific) in loading buffer composed of water and 0.1% formic acid. Peptides were eluted with a Mobile phase B gradient from 2% to 35% B in 9.6 min. Eluting peptide cations were converted to gas-phase ions by electrospray ionization and analyzed on a Thermo Orbitrap Astral (Thermo Scientific) by data independent approach. Survey scans of peptide precursors from 380 to 980 m/z were performed in orbitrap at 120K resolution (at 200 m/z) with a 5 × 10^6^ ion count target. DIA scans were performed in astral analyzer. AGC target was set to 5 × 10^4^ and maximum injection time to 2 ms. Precursor mass range 380 – 980 m/z was covered by 4 m/z windows. Activation type was set to HCD with 25% collision energy. All data were analyzed and quantified with the Spectronaut 19 software [72] using direct DIA analysis. A spectral library was generated based on the predicted protein sequences for *T. brucei* TREU927 sourced from TriTrypDB (version 64) [73]. Enzyme specificity was set as C-terminal to Arg and Lys, also allowing cleavage at proline bonds and a maximum of two missed cleavages. Carbamidomethylation of cysteines was set as fixed modification and N-terminal protein acetylation and methionine oxidation as variable modifications. FDR was set to 1 % for PSM, peptide and protein. Quantification was performed on MS2 level. Precursor PEP cutoff and precursor and protein cutoff was set to 0.01, protein PEP was set to 0.05. For generating statistical plots, *t*-test–based statistics were applied to normalized intensities and *P*-values and significance cutoffs calculated using Perseus [70].

### Bioinformatics

Sequence alignments were generated in Clustal Omega [74] and edited using Jalview [75]. Structural models were generated with Phyre2 [76] and AlphaFold 3 [77]. Molecular graphics and analyses were performed with Pymol (Version 3.1.6.1, Schrödinger, LLC).

## Acknowledgements

This work was funded by a grant from the Charles University Grant Agency (GAUK project no. 276623) to S.S., A.Z. and M.Z, by a bilateral GACR/DFG grant (project IDs.: 25-18648L and KR 4017/9-2) to M. Z. and S. K. and by a Wellcome Trust and Royal Society Sir Henry Dale Fellowship (208780/Z/17/Z) to CT. AZ is grateful for childcare support covered by a Martina Roeselova Memorial fellowship (IOCB Tech Foundation). We thank Graham Warren for supplying the anti-GRASP antibody, Jay Bangs (University at Buffalo) for rabbit anti-BiP antibodies and the OMICS Proteomics BIOCEV core facility for excellent technical service.

## Data Availability

The mass spectrometry proteomics data generated for this study have been deposited at the PRIDE repository (https://www.ebi.ac.uk/pride/) under accession codes PXD075517 (TurboID) and PXD075464 (whole cell proteomics).

## Supplementary Material

**Figure S1. Localisation of TurboID fusion proteins in *BSF T. brucei*.** (A) In *T. brucei* BSF, biotin labelling by BiP-TurboID fusion and BiPN-TurboID was detected with Cy5-streptavidin (red). The signal largely colocalises with that of anti-BiP antibody (green; Alexa488) consistent with an ER stain. Shown are single plane raw images for DAPI (blue), anti-BiP, Cy5-streptavidin fluorescence and a respective merge. Scale bar = 2 μm. (B) Biotinylation elicited by BiPN-TurboID proximity labelling in BSF colocalizes with the Golgi apparatus in addition to the ER, as monitored by co-staining with Cy5-streptavidin (red) and anti-GRASP (green; Alexa488). Scale bar = 5 μm. Respective localisation of BiP-TurboID fusion and BiPN-TurboID in PCF are shown in Figure 2.

**Figure S2. Localisation of TurboID fusion proteins in PCF *T. brucei* in relation to Bodipy TR Ceramide / GRASP.** (A) In *T. brucei* PCF, biotin labelling by BiP-TurboID fusion was detected with Cy3-streptavidin (magenta). The signal largely colocalises with that of anti-BIP antibody (green; Alexa488) consistent with an ER stain but not with the signal of Bodipy TR ceramide (red) which integrates preferentially into the ceramide-rich membranes of the TGN [17]. Shown are single plane raw images for DAPI (blue), anti-BiP, Cy3-streptavidin fluorescence, Bodipy TR ceramide and respective merges. (B) In contrast we observed colocalization of the streptavidin signal with an antibody signal of the cis-Golgi marker GRASP (Figure 2) which also colocalizes with an anti-HA antibody signal detecting the BiP- and BiPN-TurboID baits directly. Shown are single plane raw images for DAPI (blue), anti-GRASP (green), anti-HA (red) and a respective merge. Scale bar = 5 μm.

**Figure S3. Tb927.11.1440 is a potential homolog of the eukaryotic ER-resident selenoprotein Sep15.** (A) Sequence alignment of Tb927.11.1440 and *D. melanogaster* Sep15. Secondary structure elements of the Sep15 NMR solution structure (pdb:2A4H; [81], spanning Sep15 residue 62- 178 and the predicted N-terminal signal peptide are indicated. Yellow circles mark Sep15 cysteine residues (strikethrough circles indicate no matching cysteines in the alignment). (B) Cartoon representations of the Sep15 (magenta) and a Phyre2 model Tb927.11.1440 (blue) with cysteine residues drawn as yellow sticks. ***T. brucei* encodes several pERp1 homologs** which are strongly labelled by BiP-TurboID. (C) Shown is a multiple sequence alignment of Tb927.11.5180, Tb927.10.9150, Tb927.11.15210 (MSAP) and murine pERp1. Secondary structure elements of the *M. musculus* pERp1crystal structure (pdb:7AAH; [25]), spanning residue 20-185 are indicated. Yellow circles mark pERp1 cysteine residues that engage in disulfide bonds (formed between the numbered cysteine positions); additional cysteines that do not match pERp1 in any sequence are marked by asterisks. (D) Below are cartoon representations of the pERp1 structure and Phyre2 models of the three *T. brucei* proteins, with the Cys residues numbered and (E) respective Alphafold models.

**Figure S4. BiP-TurboID life-stage differences.** Volcano plot for comparison of the BiP-TurboID experiments in PCF (right side of plot) and BSF (left side of plot), generated by plotting the − log_10_*P*-value versus the *t*-test difference. Datapoints are color-coded for TrypTag localization are in dark blue for ER, light blue for Golgi, green for surface, pink for endocytic. A statistical cut-off curve is drawn in orange (FDR=0.05, s0=0.1; see Methods part). Selected data points are labelled, for complete annotation see Table S3.

**Figure S5. Endogenous tagging confirms ER localisation of EroI (Tb927.8.4890).** Anti-HA immunoblot of BSF *T. brucei* parental cells and six independent EroI-6xHA clones. A band at approximately 70 kDa, consistent with the predicted molecular weight of EroI-6xHA with seven putative *N*-glycosylation sites, expected an increase in apparent molecular weight of approximately 14 to 21 kDa, is present in 3 positive clones 4, 5, and 6 and absent from parental cells and clones 1, 2 and 3.

**Figure S6. Tb927.5.1160 is a potential homolog of the mammalian BiP NEI MANF.** Shown are cartoon representations of the crystal structure of the *Cricetulus griseus* BiP nucleotide binding domain (light gray) in complex with murine MANF (light blue for SAP domain /dark blue for SAP-like domain; compare Figure 4 A) (pdb:6HA7) (left), a respective Alphafold3 model of the same sequences but with the entire BiP sequence (including the client binding domain (dark gray)) and an Alphafold3 model of *T. brucei* BiP in complex with Tb927.5.1160 (red for SAP domain /orange for C-terminal domain; compare Figure 4 A). Alphafold3 model confidence metrics are shown at the bottom.

**Figure S7. Impact of Tb927.5.1160 silencing and inducible exogenous expression changes on cell growth and ER stress sensitivity.** Shown are growth curves (n=3; insets show population doubling time (PDT)) and dose response curves for dithiothreitol (DTT) (n>6; insets show EC_50_ values) comparing tetracycline induced cells with uninduced controls for (A) Tb927.5.1160 RNAi in PCF, (B) RNAi in BSF and (C) Tb927.5.1160 inducible expression in BSF. The western blot in (C) detects expressed Tb927.5.1160 (asterisk) via the C-terminally fused 6xmyc epitope.

**Figure S8. Impact of TbMANF silencing and inducible expression changes on the whole cell proteome.** Cells induced for 48 h (A) for RNAi depletion of Tb927.5.1160 in PCF or (B) for RNAi depletion of Tb927.5.1160 in BSF and (C) for Tb927.5.1160 overexpression were harvested and analysed for protein abundance compared to uninduced cells. Data points associated with GO-annotations or TrypTag localisation for ER or Golgi apparatus are colored in the statistical volcano plots (see inset in (A) for color code and hierarchy). Selected datapoints are labelled and corresponding proteins detailed in in Table S3. All datasets were subjected to a consistent statistical analysis with a significance cut-off (orange curve) determined applying equal criteria (FDR=0.05, s0=0.1; see Methods part). Kua, Tb927.10.2250; PT1, Tb927.3.4070; PT4, Tb927.3.4100; DnaJ_1, Tb927.2.3960; DnaJ_2, Tb927.3.1760; ERGIC, Tb927.3.1760; AAT1_1, Tb927.8.7620/30; AAT1_2, Tb927.8.7610.

**Table S1. Primers**

**Table S2. Canonical ER-resident proteins detected by BiP-TurboID Table S3. Proteomics Table TurboID**

**Table S4. Summary of surface and cargo detections**

**Table S5. Proteomics Table TbMANF RNAi and inducible expression**

**Table S6. Summary of protein abundance changes elicited by TbMANF RNAi and inducible expression**

