## Supplementary material for "A luminal proteome of the endoplasmic reticulum and Golgi apparatus reveals a novel modulator of ER stress tolerance in African trypanosomes": Table S1

| **Primers** | **sequence** | **purpose** |
| --- | --- | --- |
| **Endogenous tagging with TurboID_2HA** | | |
| BiP-2HA-TurboID-MDDL_F | AGAGCGTGACCAACCCCATCATCCAGAAGACTTACCAATCCGCCGGCGGAGGCGACAAGCCACAACCCATGGACGATCTGGGTTCTGGTAGTGGTTCC | BiP-2HA-TurboID |
| BiP-2HA-TurboID-MDDL_R | ATTTAAACGAACCAAATTTGCTGCAGTAATTATAAAAAACGGCATTATAAAACAAACGATTCAAACACCTGCCAGTACACCCAATTTGAGAGACCTGTGC | BiP-2HA-TurboID |
| NUP65-2HA-TurboID | AGCCGTTTAACGAAGTTGTACCGAACGTATTGTTGATCGACAGAGTTCATGGAAGCGACAAATACAGGGCAAAAACACTCGGTTCTGGTAGTGGTTCC | NUP65-2HA-TurboID |
| NUP65-2HA-TurboID | TTTTTGATGTTGCCAAACCTCTGTCGTAGGAAACTGGACATATAACACACCACCACAAATTGTTCTTCTTGGGTAAAATACCAATTTGAGAGACCTGTGC | NUP65-2HA-TurboID |
| **overexpression** | | |
| BiPN-TurboID-HA_F | GCGCAAGCTTCCGCCGCCATGTCGAGGATGTGGCTGACCAC | Expression of BiPN-TurboID-HA |
| BiPN-TurboID-HA_R | GCGCGGATCCCCCGCCAACCTCGCTTTCACCG | Expression of BiPN-TurboID-HA |
| TbMANFOE_F | GCGCAAGCTTATGTTCTCTATGTCGCCATGCG | TbMANF Expression  via pRPAtag |
| TbMANFOE_R | GCGCAAGCTTATGTTCTCTATGTCGCCATGCG | TbMANF Expression  via pRPAtag |
| **Stem-loop RNAi** | | |
| TbMANF_F | ATTTAAATCAGCCGTCCTTGTGGTTTTG | TbMANF_RNAi  In PCF via p3666 |
| TbMANF_R | GGATCCGGAAGTCTTGAGAAGCGCCT | TbMANF_RNAi  In PCF via p3666 |
| TbMANF_RNAi_Bsp120I_Acc65i_F | GATCGGGCCCGGTACCCAGCCGTCCTTGTGGTTTTG | TbMANF_RNAi  In BSF via pRPAISL |
| TbMANF_RNAi _XbaI_BamH1_R | GATCTCTAGAGGATCCGGAAGTCTTGAGAAGCGCCT | TbMANF_RNAi  In BSF via pRPAISL |
| LEM3_F | ATTTAAATGGAGGCCTCCTTTCGATACG | RNAi  In PCF via p3666 |
| LEM3_R | GGATCCACCGTTCTTGTCGCAGTGAT | RNAi  In PCF via p3666 |
| Tb927.10.3430_F | ATTTAAATACCGCGGTAGTAACTTGTGG | RNAi  In PCF via p3666 |
| Tb927.10.3430_R | GGATCCCGCAGTAATGCCTCGAAAGC | RNAi  In PCF via p3666 |

| **CRISPR Cas9 endogenous C-terminal tagging primers** | |
| --- | --- |
| **EroI (Tb927.8.4890)** | |
| downstreamF | TGCGAAAAATTCAATTACACGGCTATGAGTggttctggtagtggttccgg |
| downstream R | AGGCTTTAAATTGAACGTCCTACCCCACCAccaatttgagagacctgtgc |
| 3'sgRNA | gaaattaatacgactcactataggATTGTCTGCTTGCACAAAGTgttttagagctagaaatagc |
| **TbMANF (Tb927.5.1160)** | Sequence 5’ to 3’ |
| downstreamF | TACGAGGTGCTAGACGGCCGTGGCGATCTTggttctggtagtggttccgg |
| downstream R | CCTCCTTTCGCCCACATTTACCTCACACGGccaatttgagagacctgtgc |
| 3'sgRNA | gaaattaatacgactcactataggGCACGAGCAAGGACAGAATAgttttagagctagaaatagc |
