## Supplementary material for "A luminal proteome of the endoplasmic reticulum and Golgi apparatus reveals a novel modulator of ER stress tolerance in African trypanosomes": Table S2

| Gene ID | Description | Difference  BIP vs wt  PCF | Difference  BIP vs wt  BSF | Ratio  BIPN/BIP  PCF | Ratio  BIPN/BIP  BSF | Localisation & Comments |
| --- | --- | --- | --- | --- | --- | --- |
| **Endoplasmic reticulum** | | | | | | |
| Tb927.9.4500 | Hsp70.a | 10.8 | 12.9 | 0.63 | 0.76 | **LOPIT (PCF&BSF):** ER  **TrypTag N&C-term:** ER |
| Tb927.9.9860 | GRP170 (Hsp70-like) | 13.1 | 14.6 | 0.73 | 0.59 | Potential BIP NEF  (PMID: 30710085)  **LOPIT (PCF&BSF):** ER  **TrypTag C-term:** cytosolic (points) |
| Tb927.11.11330 | Hsp70.4 | 6.5 | 0.8 | 2.74 | 1.21 | **LOPIT (PCF&BSF):** cytosol  **TrypTag N&C-term:** cytoplasm(patchy), flagellar cytoplasm |
| Tb927.3.3580 | GRP94/LPG3(Hsp90) | 13.3 | 16.6 | 2.45 | 1.16 | **LOPIT (PCF&BSF):** ER  **TrypTag C-term:**  ER, Golgi |
| Tb927.10.10980* | Hsp90 | 4.1 | 1.5 | 2.62 | 0.89 | **TrypTag N-term:**  cytoplasm, nucleoplasm |
| Tb927.3.1430 | DnaJB11 | 7.1 | 9.0 | 0.32 | 0.26 | ER (PMID: 30506377) |
| Tb927.11.16740 | DnaJ | 7.5 | 7.9 | 1.03 | 1.68 | **LOPIT (PCF&BSF):** ER  **TrypTag C-term:**  ER, Golgi |
| Tb927.11.5220 | DnaJ | 7.7 | 2.1 | 0.33 | 0.78 | **LOPIT (PCF&BSF):** ER  **TrypTag C-term:** ER, nuclear envelope |
| Tb927.2.5160 | DnaJ2 | ND | 1.1 | ND | 0.81 | **LOPIT (PCF&BSF):** cytosol  **TrypTag N-term:** cytoplasm (points), endocytic**; C-term:** cytoplasm, flagellar cytoplasm, nucleoplasm |
| Tb927.7.3630 | DnaJ (TPR-repeat) | 8.2 | 11.7 | 0.43 | 0.40 | **LOPIT (PCF&BSF):** ER  **TrypTag C-term:** cytoplasm(reticulated, weak) |
| Tb927.9.10010 | Sec63 | ND | 7.2 | ND | BIP only | (PMID: 18768469)  **LOPIT (PCF&BSF):** ER |
| Tb927.5.1160 | Degradation arginine-rich protein for mis-folding | 13.3 | 17.4 | 1.14 | 0.72 | Potential BiP interactor (NEI)  ER (this work) |
| Tb927.11.5180 | Potential pERp1 homolog | 14.0 | 16.9 | 0.80 | 0.90 | **LOPIT (PCF&BSF):** ER  **TrypTag C-term:** ER, endocytic |
| Tb927.11.15210 | TLR4 regulator and MIR-interacting MSAP; Esp16 | 12.0 | 15.3 | 1.0 | 0.8 | **LOPIT (PCF&BSF):** ER  **TrypTag C-term:** cytoplasm (reticulated)  PMID: 25931509 |
| Tb927.8.7410 | Calreticulin | 12.2 | 14.9 | 0.98 | 0.98 | **LOPIT (PCF&BSF):** ER |
| Tb927.10.13630 | ER glucosidase II | 10.9 | 11.6 | 0.67 | 0.60 | **LOPIT (PCF&BSF):** ER  **TrypTag C-term:** cytoplasm (weak, reticulated) |
| Tb927.3.4630 | UGGT (UDP-glucose:glycoprotein glucosyltransferase) | 11.3 | 14.4 | 0.76 | 0.74 | **LOPIT (PCF&BSF):** ER  **TrypTag N-term:** cytoplasm (weak, reticulated)**; C-term:** ER, cytoplasm (weak, reticulated) |
| Tb927.11.10700 | Yos9 | 8.2 | 9.5 | 1.36 | 0.84 | **LOPIT (PCF&BSF):** ER  **TrypTag C-term:** endocytic, cytoplasm(points) |
| Tb927.11.4200 | ERGIC53 | 7.0 | 3.5 | 2.43 | 0.78* | **LOPIT (PCF&BSF):** secretory endocytic 1  **TrypTag N-term:** cytoplasm, flagellar cytoplasm, nuclear lumen |
| Tb927.4.2450 | PDI1 | 11.4 | 13.9 | 0.32 | 0.34 | **LOPIT (PCF&BSF):** ER  **TrypTag N-term:** cytoplasm (points); **C-term:** cytoplasm (weak, reticulated, points) |
| Tb927.7.1300 | PDI (ERp72-like) | 10.0 | 12.4 | 1.29 | 0.83 | **LOPIT (PCF&BSF):** ER  **TrypTag C-term:**  ER, Golgi |
| Tb927.10.8230 | PDI2 (ERp57-like) | 9.9 | 13.3 | 0.80 | 1.26 | **TrypTag N&C-term:** endocytic, cytoplasm (reticulated, weak) |
| Tb927.7.5790 | PDI (Erp44-like) | ND | 8.8 | ND | 0.50 | **LOPIT (PCF&BSF):** ER |
| Tb927.5.1020 | PDI | 10.8 | 11.7 | 1.13 | 0.81 | **LOPIT (PCF&BSF):** ER  **TrypTag N&C-term:** endocytic, cytoplasm (reticulated, weak) |
| Tb927.8.4890 | Ero1 | 11.3 | 13.0 | 1.40 | 1.17 | ER (this work) |
| Tb927.11.6230 | Sec61 | ND | ND | ND | Only BIPN | **LOPIT (PCF&BSF):** ER  **TrypTag N-term:** ER |
| Tb927.5.1930 | Signal peptidase subunit SPC3, putative | 10.7 | 13.7 | 2.0 | 0.9 | **LOPIT (PCF&BSF):** ER  **TrypTag N&C-term:** ER, nuclear envelope (PMID: 25816352) |
| Tb927.5.3220 | signal peptidase type I, putative | borderline | 7.2 | ND | 0.39 | **LOPIT (PCF&BSF):** ER  **TrypTag N-term:** ER; **C-term:** ER, nuclear envelope (strong) |
| Tb927.8.2910 | EDEM1 | 8.8 | 9.1 | 1.2 | 0.8 | **TrypTag C-term:** ER (75%), cytoplasm (reticulated, weak) |
| Tb927.8.2940; Tb927.8.2930 | EDEM4; EDEM3 | 7.5 | ND | 0.92 | Only BIPN borderline detection | **TrypTag C-term:** cytoplasm (reticulated); - |
| Tb927.8.2920 | EDEM2 | 6.4 | 1.0 | Only detected in BIP | Only BIP borderline detection | - |
| Tb927.4.590 | EMC1 | 9.6 | 10.0 | 0.4 | 0.5 | **LOPIT (PCF&BSF):** ER  **TrypTag C-term:** ER, nuclear envelope  (PMID: 35500022) |
| Tb927.10.13290 | Ethanolamine phosphotransferase | ND | 4.0 | ND | Only detected in BIP | **LOPIT (PCF&BSF):** ER  **TrypTag N-term:** mitochondrion; **C-term:** mitochondrion, nuclear envelope |
| Tb927.10.13860 | GPI transamidase subunit 8 (GPI8) | 9.0 | 11.0 | 1.6 | 1.0 | **LOPIT (PCF&BSF):** ER  **TrypTag C-term:** ER, nuclear envelope |
| Tb927.4.1920 | GPI transamidase subunit 16 (GPI6) | 9.2 | 11.2 | 2.0 | 0.9 | **LOPIT (PCF&BSF):** ER  **TrypTag N-term:** endocytic, cytoplasm(weak) |
| Tb927.11.15760 | GPI transamidase subunit Tta1 (Gab1) | 7.9 | 10.0 | 1.8 | 1.7 | Robustly detected in both stages |
| Tb11.v5.0339 | GPI transamidase component Tta2 | 6.2 | 7.7 | 0.6 | 0.5 | - |
| Tb927.10.210 | GPI transamidase component GAA1 | 1.9 | 7.4 | 2.9 | 0.6 | **LOPIT (PCF&BSF):** ER  **TrypTag N-term:** ER |
| Tb927.5.4600 | GRESAG3 | ND | 9.9 | ND | 0.7 | ER (this work) |

*Tb927.10.10980;Tb927.10.10960;Tb927.10.10940;Tb927.10.10910;Tb927.10.10970;Tb927.10.10950;Tb927.10.10920;Tb927.10.10900;Tb927.10.10930;Tb927.10.10890;Tb11.v5.0543

**Table S2. Overview of known luminal protein components of the *T. brucei* secretory pathway**
