## Supplementary material for "A luminal proteome of the endoplasmic reticulum and Golgi apparatus reveals a novel modulator of ER stress tolerance in African trypanosomes": Table S4

| Gene ID | Description | Difference  BIP vs wt  PCF | Difference  BIP vs wt  BSF | Ratio  BIPN/BIP  PCF | Ratio  BIPN/BIP  BSF | log_2_-fold  Difference  PCF/BSF  (PMID: 26910529) |
| --- | --- | --- | --- | --- | --- | --- |
| **Surface proteins** | | | | | | |
| Tb927.5.390 | ISG75 | ND | 9.1 | ND | 0.8 | -4.5 |
| Tb927.5.360 | ISG75 | 2.5 | 9.8 | 2.3 | 1.3 | -5.4 |
| Tb927.2.3270 | ISG65 | ND | ND | ND | Inf | -4.2 |
| Tb927.2.3320 | ISG65 | ND | ND | ND | Inf | -4.0 |
| Tb927.1.4900 | ESAG11 | ND | 4.9 | ND | 1.9 | -4.9 |
| Tb927.7.3260;  Tb927.7.3250 | ESAG6/ESAG7 | ND | 9.5 | ND | BiP only | -6.0 |
| Tb927.5.120 | ESAG9 | ND | 8.3 | ND | 2.3 | -4.5 |
| Tb927.1.5100 | ESAG2 | ND | 10.2 | ND | 1.2 | -3.4 |
| Tb927.3.570 | ESAG2 | ND | 7.4 | ND | BiP only | 0.1 |
| Tb927.5.4600;  Tb11.v5.0367;  Tb05.5K5.240 | ESAG3 | ND | 9.9 | ND | 0.7 | NaN |
| Tb11.v5.0752 | ESAG3 | ND | 11.5 | ND | 0.6 | NaN |
| Tb927.5.340 | ESAG5 | ND | ND | (BiPN only) | ND | -4.5 |
| Tb927.7.6860 | ESAG5 | ND | 8.4 | ND | 0.6 | 2.7 |
| Tb11.v5.0402* | BARP | borderline | 3.3 | 3.7 | 4.7 | NaN |
| Tb927.9.15590 | BARP | ND | borderline | ND | 13.6 | NaN |
| Tb927.10.5680 | PAG1 | ND | 9.1 | ND | 2.5 | NaN |
| Tb927.10.5690 | PAG2 | ND | 9.8 | ND | 2.0 | -4.0 |
| Tb927.10.10240 | PAG1 | 5.8 | ND | 1.3 | ND | NaN |
| Tb927.10.10220 | PAG2* | ND | ND | (BiPN only) | ND | NaN |
| Tb927.11.14610 | PAG4 | ND | 6.2 | ND | 2.0 | -5.2 |
| Tb927.10.10230 | PAG5 | 7.1 | ND | 1.1 | ND | NaN |
| Tb927.7.6850 | Trans-sialidase | 8.8 | ND | 0.8 | ND | 3.8 |
| Tb927.8.7350 | Trans-sialidase | 9.0 | ND | 0.5 | ND | NaN |
| Tb927.8.7340 | Trans-sialidase | 10.0 | ND | 0.3 | ND | NaN |
| Tb927.11.11410 | Trans-sialidase | ND | 7.3 | ND | 0.9 | 1.5 |
| Tb927.7.4260 | ESP13 | ND | ND | ND | Inf | -3.5 |
| Tb927.8.1630;  Tb927.8.1620;  Tb927.8.1610 | MSP-B major surface protease | 10.6 | ND | 1.1 | ND | 3.2 |
| Tb927.8.1640 | MSP-B major surface protease | 2.1 | ND | (BiP only) | ND | 3.2 |
| Tb927.8.1610; Tb927.8.1620; Tb927.8.1630 | GP63 | 10.6 | ND | 1.1 | ND | 3.18 |
| Tb927.11.7710 | Gp63 | ND | 9.8 | ND | 0.2 | NaN |
| Tb927.11.7720 | Gp63 | ND | 9.2 | ND | BiP only | NaN |
| Tb11.v5.0710 | Gp63 | ND | 8.4 | ND | BiP only | NaN |
| Tb927.1.5170 | VSG-related | ND | 3.9 | ND | BiP only | NaN |
| Tb927.5.291b | VSG | 5.6 | 12.8 | 0.5 | 0.7 | NaN |
| Tb927.8.4010 | FLA1 | 7.0 | ND | 0.6 | ND | 3.6 |
| **Cargo** | | | | | | |
| Tb927.5.4060 | Nicastrin | 6.1 | ND | 0.3 | (BiPN only) | 0.1 |
| Tb927.4.2500 | eIF2K2 | 8.5 | 10.1 | 1.7 | 1.2 | -0.4 |
| Tb927.3.3450 | Arl3A | ND | 7.4 | (BiPN only) | 1.0 | -1.7 |
| Tb927.6.3650 | Arl3C | borderline | 11.3 | 0.7 | 0.8 | -0.3 |
| Tb927.6.990** | CatL | 3.3 | 8.5 | 2.4 | 0.4 | -1.3 |
| Tb927.11.11520 | PEX11 | 6.6 | 10.7 | 0.2 | 0.6 | 0.2 |
| Tb927.9.11580 | Gim5A | 5.9 | 10.3 | 0.1 | 0.2 | NaN |

ND=not detected; borderline=low confidence detection; BiPN/BiP ratios in brackets are based on low confidence detections

*Tb11.v5.0402;Tb11.v5.0343;Tb927.9.15580;Tb927.9.15610;Tb927.9.15630;Tb927.9.15540;Tb11.v5.0346;Tb927.9.15510;Tb11.v5.0403

**Tb927.6.990;Tb927.6.1060;Tb927.6.1050;Tb927.6.1000;Tb927.6.980;Tb927.6.970;Tb927.6.960;Tb927.6.1020;Tb927.6.1010;Tb927.6.1040;Tb927.6.1030

**Table S4. Overview of known surface and cargo proteins**
