## Supplementary figures and images for "A luminal proteome of the endoplasmic reticulum and Golgi apparatus reveals a novel modulator of ER stress tolerance in African trypanosomes"

### Figure S1

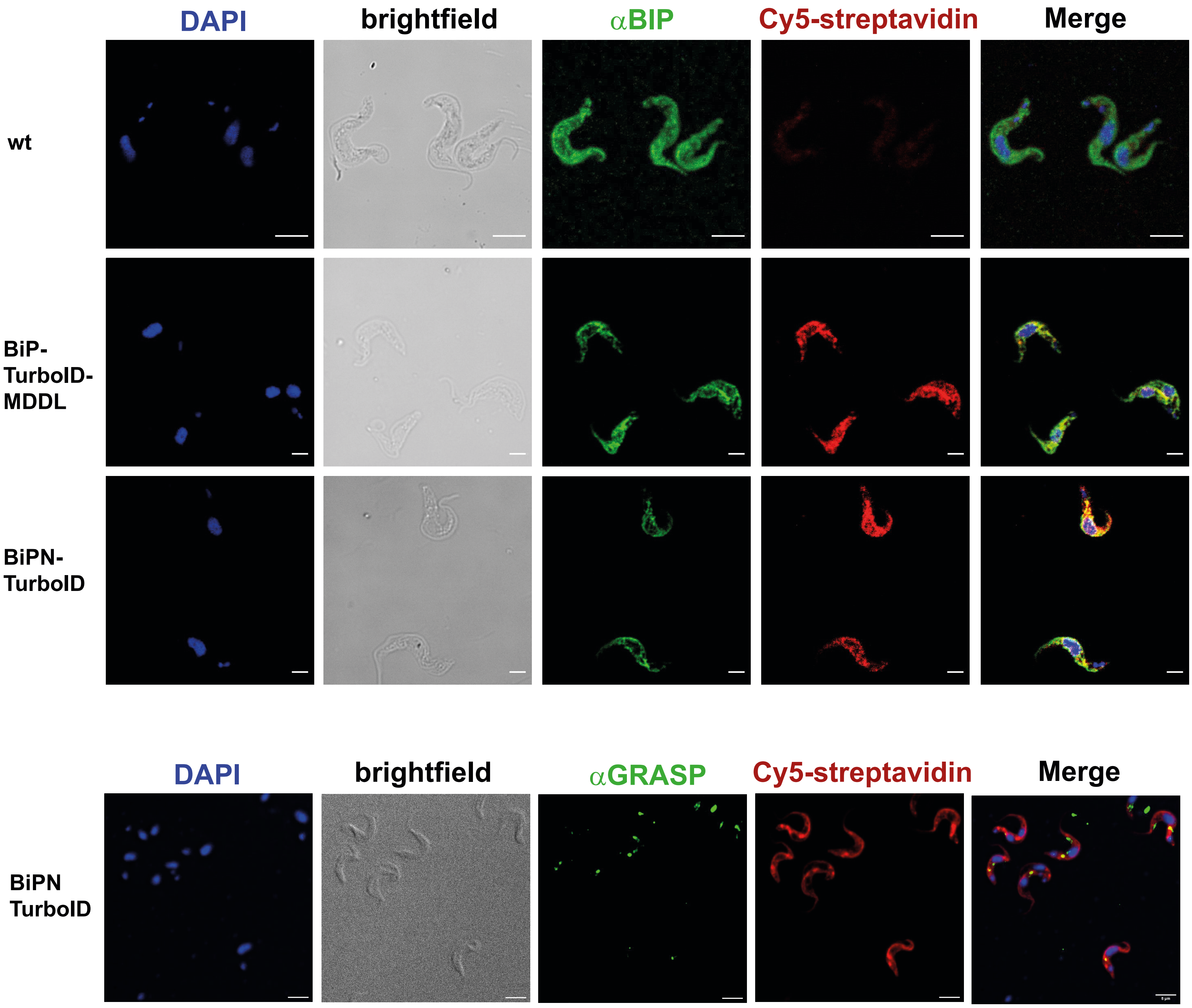

### Figure S2

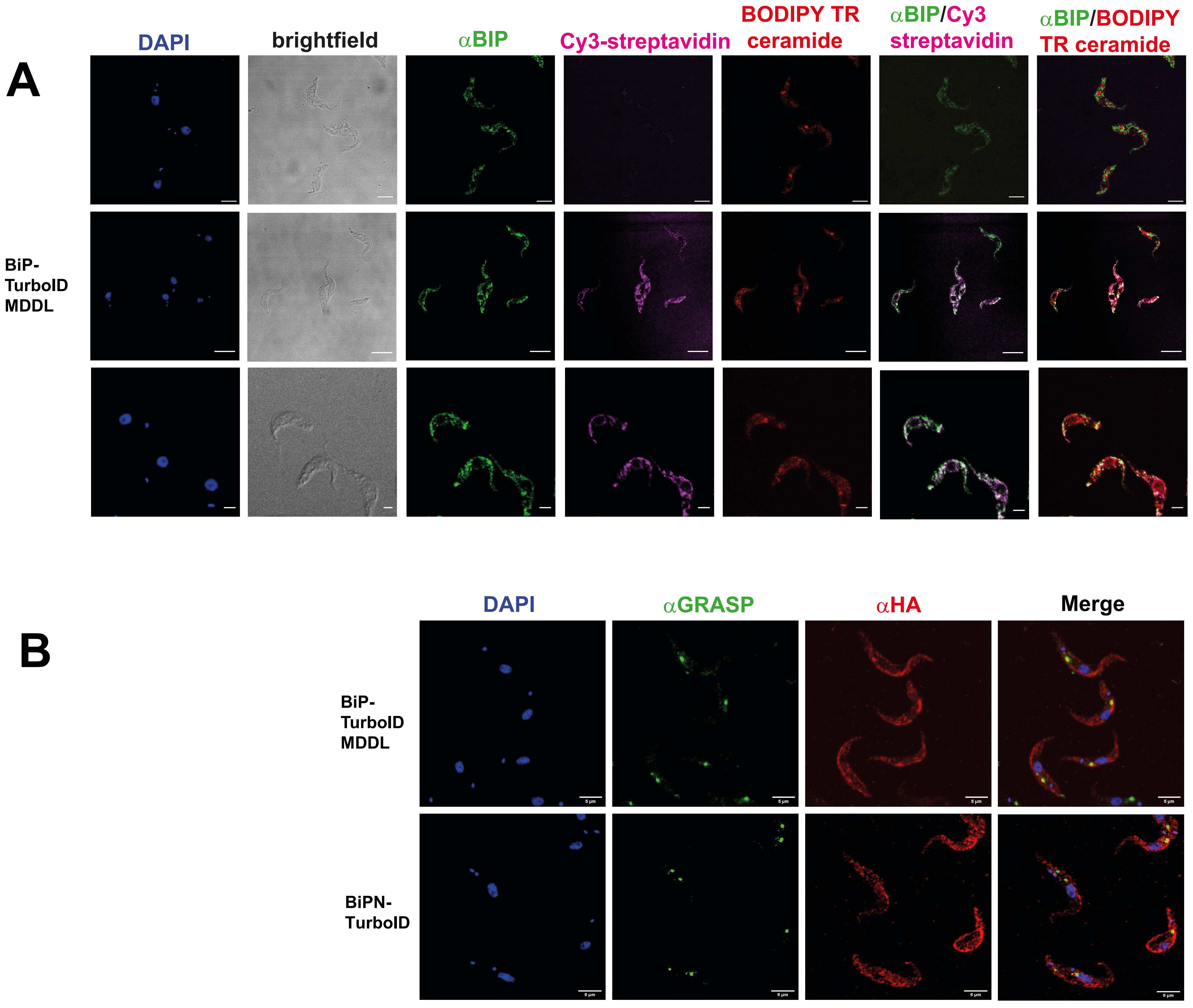

### Figure S3

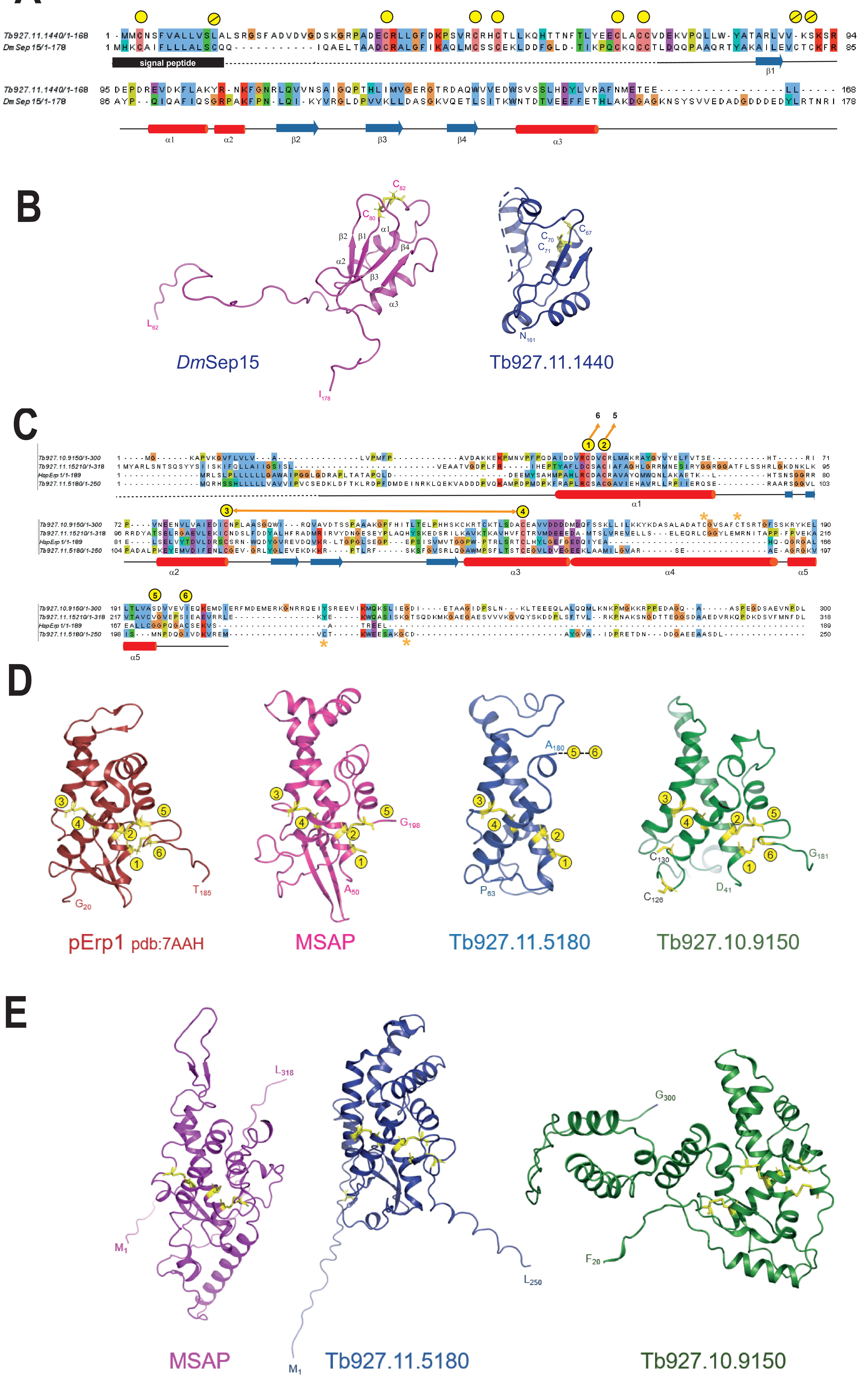

### Figure S4

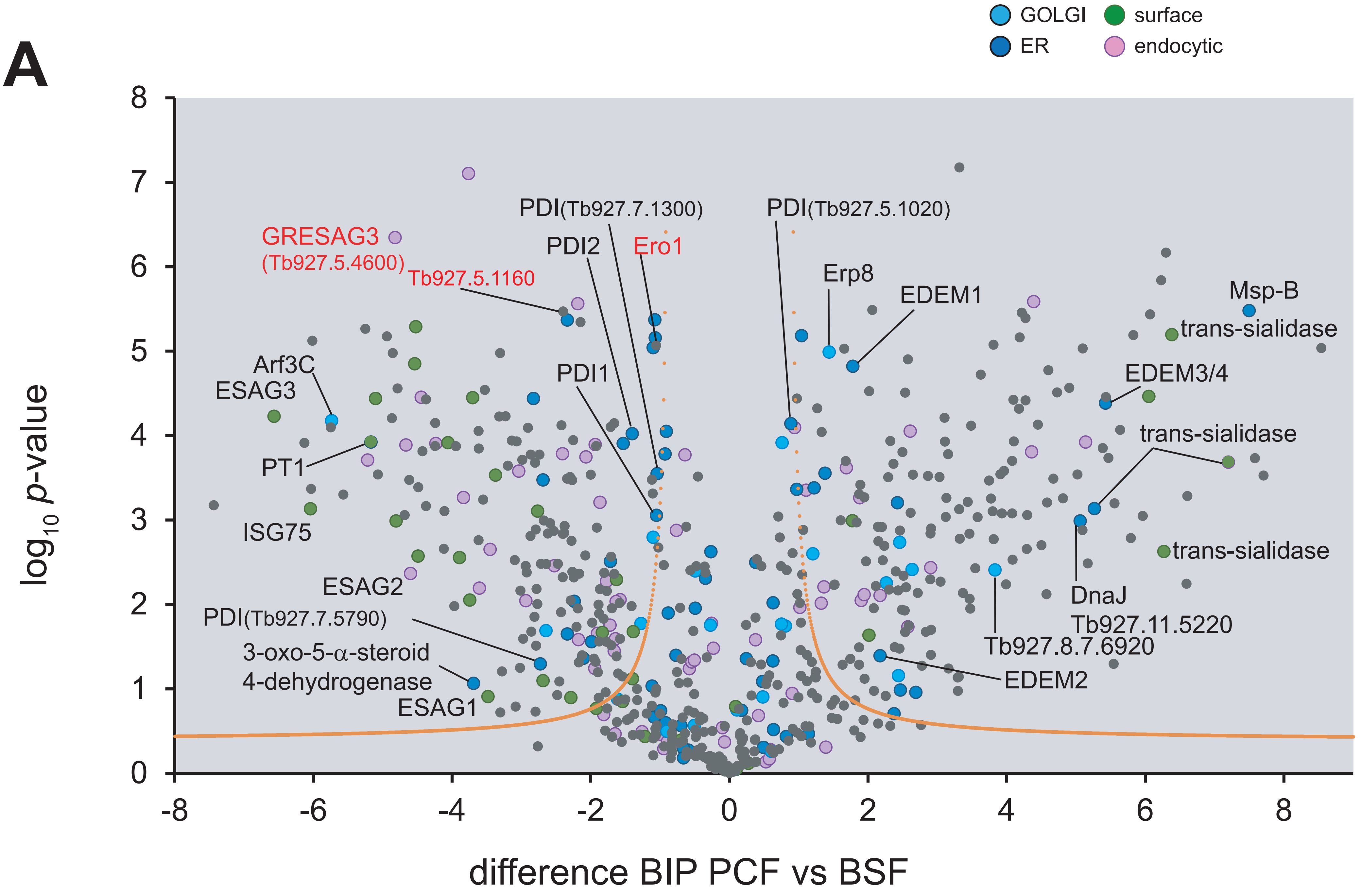

### Figure S5

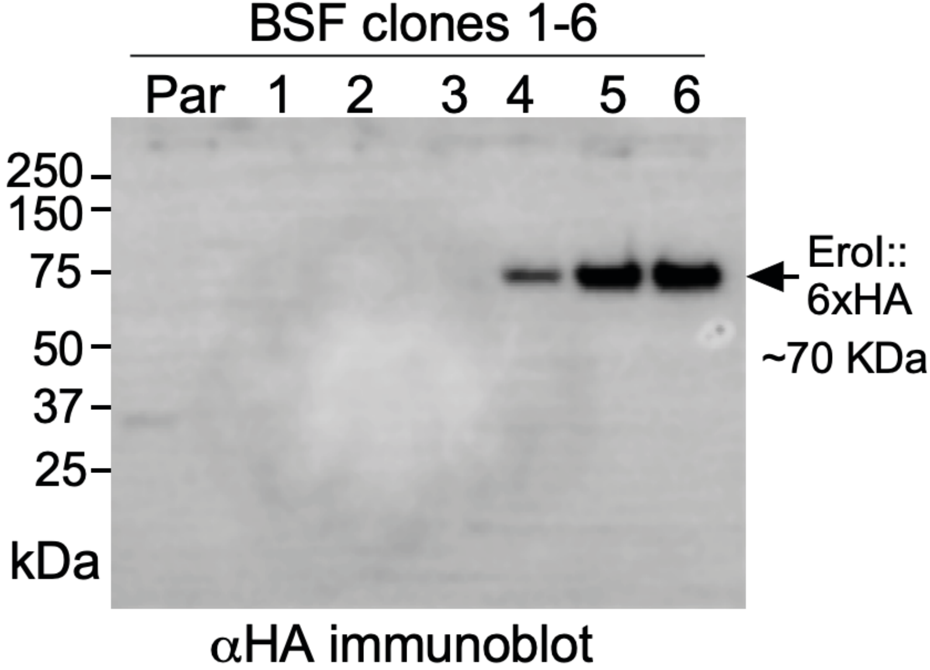

### Figure S6

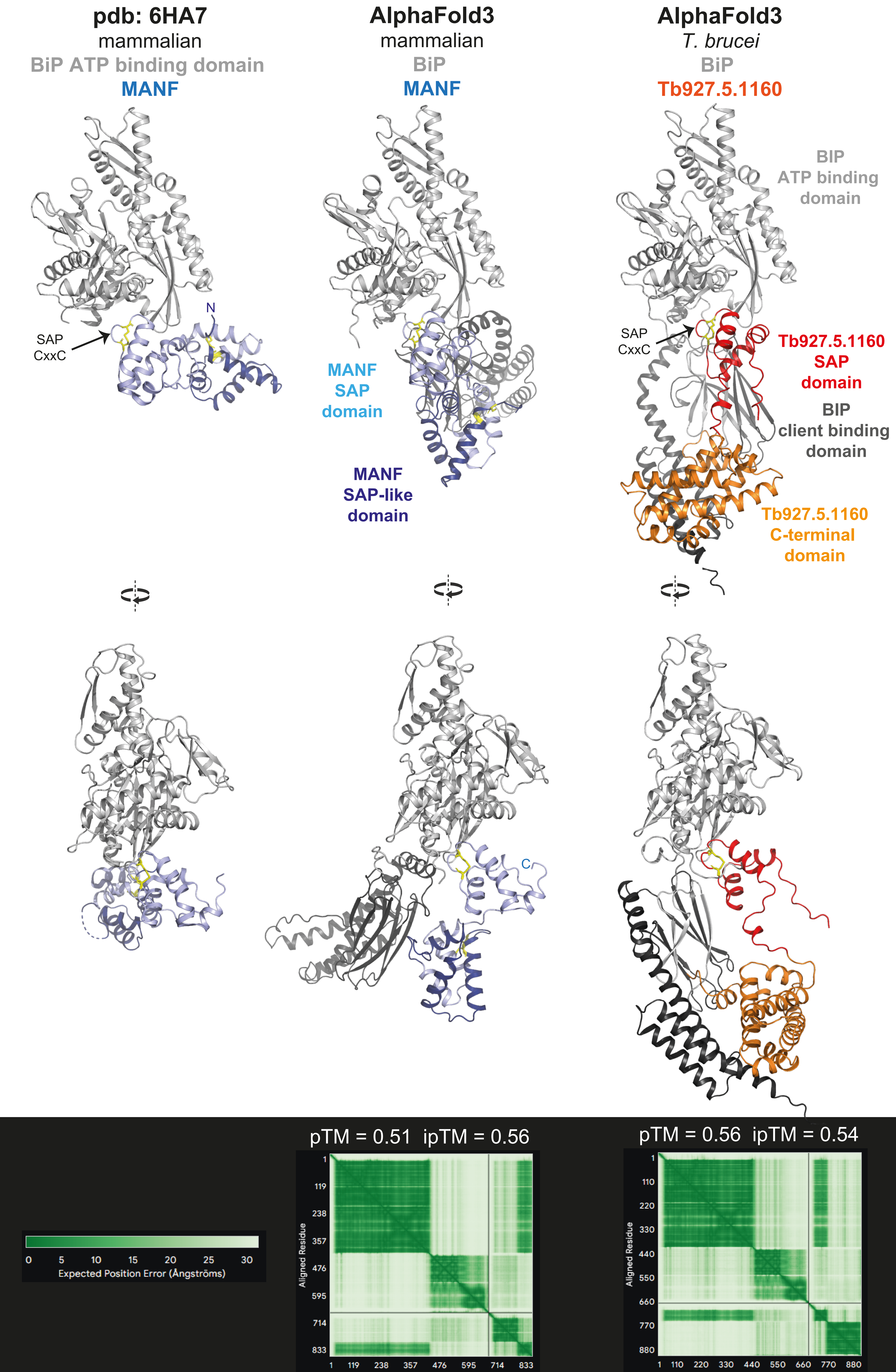

### Figure S7

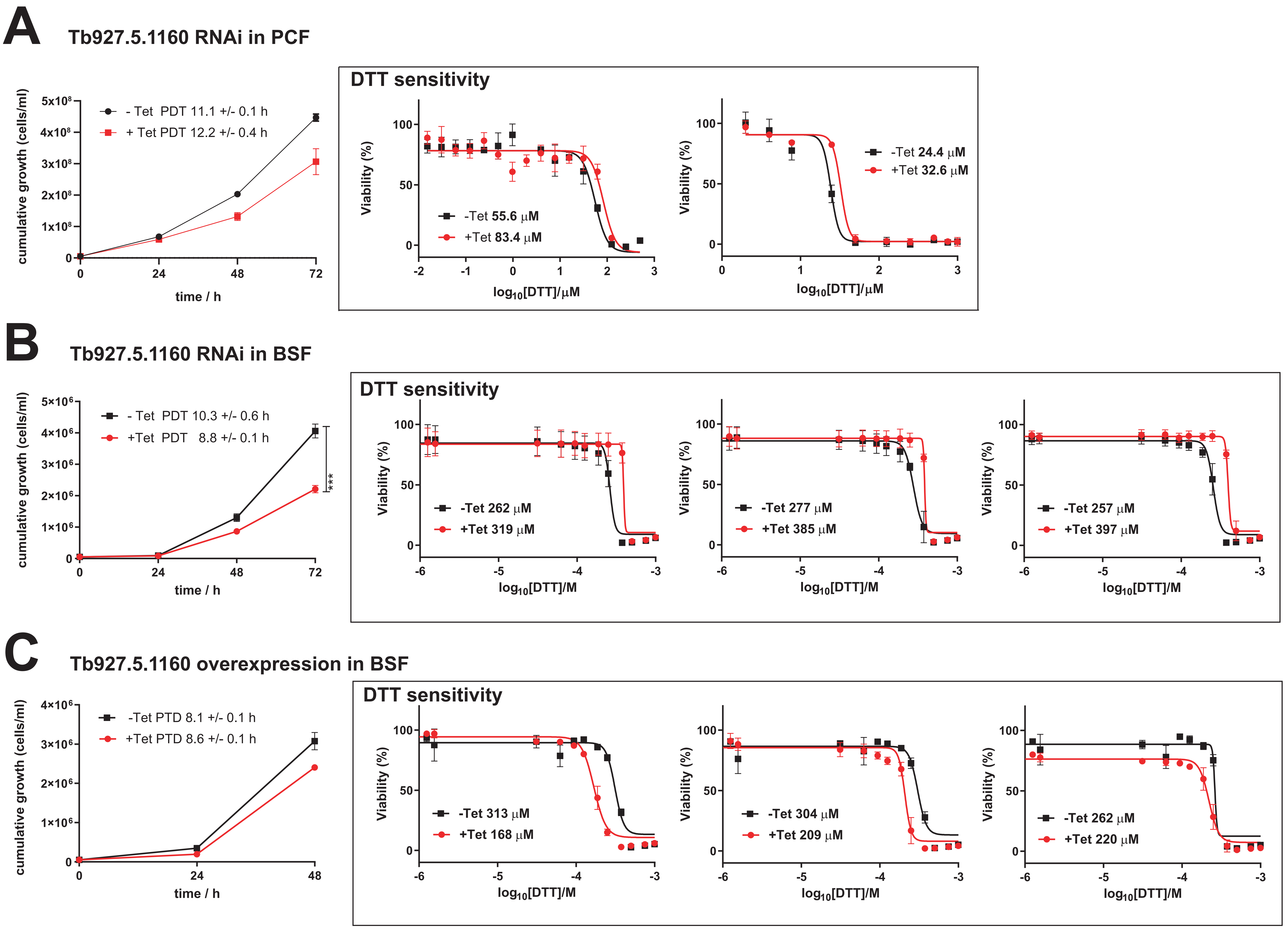

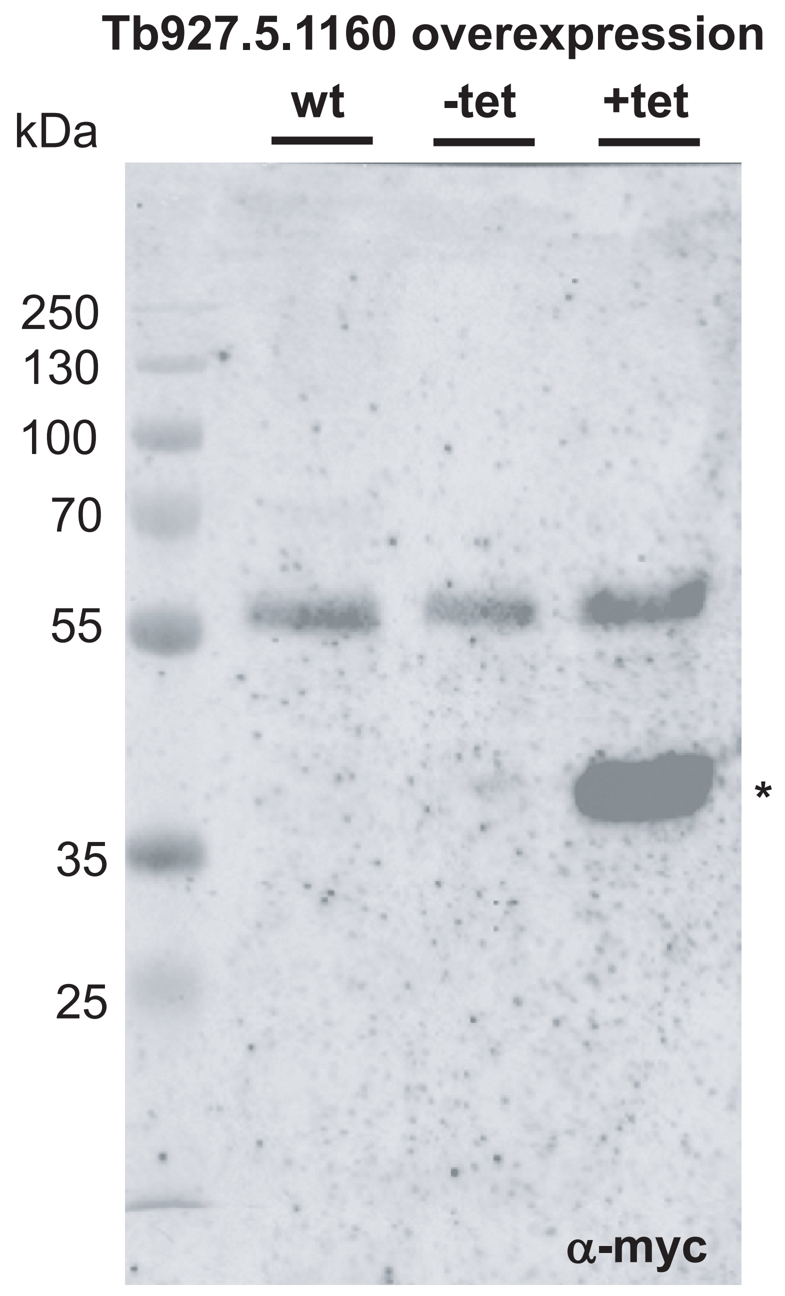

### Figure S8

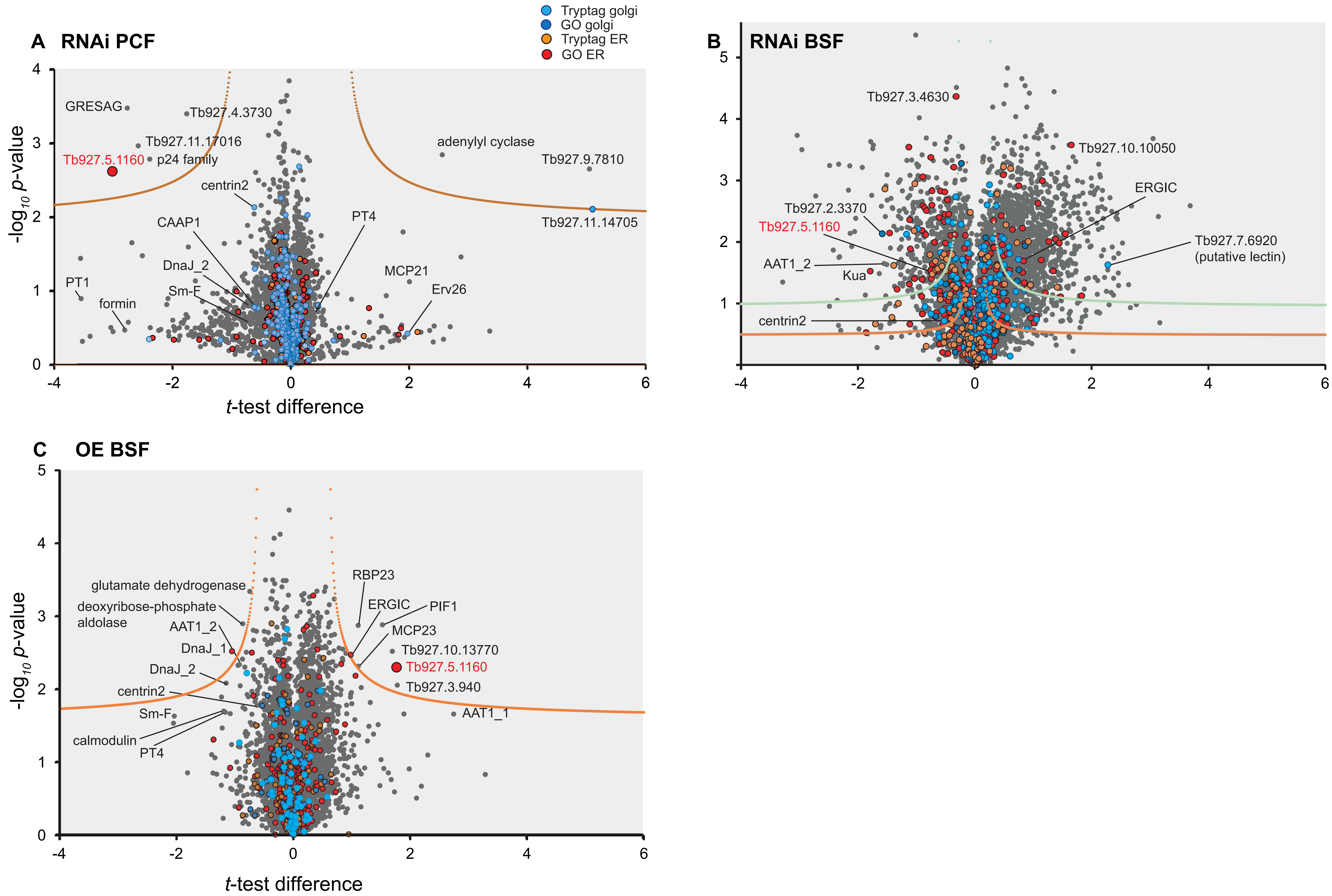
